# The impact of FreeSurfer versions on structural neuroimaging analyses of Parkinson’s disease

**DOI:** 10.1101/2024.11.11.623071

**Authors:** Andrzej Sokołowski, Nikhil Bhagwat, Dimitrios Kirbizakis, Yohan Chatelain, Mathieu Dugré, Jean-Baptiste Poline, Madeleine Sharp, Tristan Glatard

## Abstract

Image processing software impacts the quantification of brain measures, playing an important role in the search for clinical biomarkers. We investigated the impact of the variability between FreeSurfer releases on the estimation of structural brain measures in Parkinson’s disease (PD). Structural brain scans from 106 controls and 209 patients were analyzed with FreeSurfer versions 5.3, 6.0.1, and 7.3.2, including longitudinal data from 125 patients. First, we measured the differences in the estimation of volume, surface area, and cortical thickness between FreeSurfer versions. Second, we focused on the relationship between MRI-derived brain measures and group differences as well as disease severity clinical outcomes, which were evaluated both cross-sectionally and longitudinally. We found high software-induced variability in the estimation of all three structural measures, which impacted clinical outcomes. There were differences between software versions in group differences between patients and healthy controls in subcortical volume and vertex-wise cortical thickness. Software variability also impacted the estimated relationship between brain structure and disease severity in patients. Hence, software variability not only relates to the estimation of structural measures, but it also impacts clinically-relevant MRI measures. Our study provides insight into the reproducibility of structural neuroimaging studies in PD populations.

## 1. Introduction

Neuroscience relies on numerous computational methods to analyze neuroimaging data. In recent years, however, the reproducibility of neuroimaging studies has been challenged. Reproducibility is characterized by two features, namely ability to replicate findings using the same analyses, and generalizability of results obtained with various data and computational approaches (Ghosh et al., 2017). The extent of the reproducibility issue in neuroimaging was most saliently shown in fMRI analysis by Botvinik-Nezer et al. (2020) who analyzed results obtained from 70 independent research teams that analyzed the same fMRI data with different processing pipelines. Teams agreed only on four out of the nine a priori hypotheses with agreement levels ranging from 21% to 37% (Botvinik-Nezer et al., 2020). One of the main factors impacting the reproducibility of neuroimaging studies is software variability (Niso et al., 2022). Notably, Bowring et al. (2019) investigated the impact of different software (AFNI, FSL, and SPM) on fMRI results. Different software pipelines produced various activation regions with overlap often lower than 50% measured with Sørensen–Dice coefficient scores, with differences in t-statistics reaching four. Aforementioned results highlight the challenges with reproducibility.

Several studies have also focused on variability of structural MRI measures, showing that software toolbox selection impacts the estimation of brain volume (Gomez-Ramirez et al., 2022; Rane et al., 2017) and cortical thickness (Bhagwat et al., 2021; Masouleh et al., 2020). Gronenschild et al. (2012) investigated the impact of FreeSurfer version on anatomical volume and cortical thickness estimation. They tested the variability between three FreeSurfer versions (4.3.1, 4.5, and 5.0) and reported significant differences in the volume estimation of numerous brain regions, both cortical and subcortical. They also found differences in the estimation of cortical thickness. Haddad et al. (2023) investigated the compatibility (measured as intraclass correlation coefficients) of brain structure estimation between three FreeSurfer versions (5.3, 6.0, and 7.1). They reported variable compatibility of cortical thickness and surface area ranging from poor to excellent depending on the brain region. There was better compatibility between the older versions (i.e., 6.0 and 5.3) than between the more recent release (i.e., 7.1) and the older ones. Another study reported relatively low correlation (r = .39 - .52) between cortical thickness measured by various software (FreeSurfer, CIVET, and ANTs) and higher (r = .83 - .89) correlation between FreeSurfer versions (5.0, 5.3, and 6.0; Bhagwat et al., 2021). Finally, Gomez-Ramirez et al. (2022) reported significant differences in subcortical volumes estimation between FreeSurfer and FSL. Prior findings justify the need to further investigate differences between software versions.

Though it is increasingly well-established that both within- and between-software differences result in brain structure measurement differences, surprisingly little research has been done to quantify the impact of these measurement differences on the relationships between brain structure and behavior. This is potentially especially important in the context of clinical neuroscience research where MRI-derived structural brain measures are often related to clinical outcomes such as neurological symptom severity or clinical disease progression. Neuroimaging has been proposed as a diagnostic and prognostic marker of neurodegenerative diseases. In patients with Parkinson’s disease (PD), MRI-derived measures such as gray matter volume and cortical thickness has been shown to be associated with disease severity (Hanganu et al., 2014; Le et al., 2020; Sarasso et al., 2021), disease progression (Gaurav et al., 2021; Tessa et al., 2014), and to differentiate between various syndromes (Chougar et al., 2021). However, currently there are no established MRI measures for the diagnosis or tracking of Parkinson’s disease, which may at least in part be due to the measurement variability and its impact on advancing our understanding of brain diseases.

The goal of this study was to investigate the impact of software variability on (1) the estimation of structural MRI-derived measures, and (2) commonly measured associations between MRI-derived brain structure and PD clinical outcomes. To our knowledge, this is the first study that investigates the impact of software variability on structural MRI measures in PD. First, we compared results obtained from three major FreeSurfer releases (i.e., 5.3, 6.0.1, and 7.3.2). We focused on the FreeSurfer toolbox as it is publicly available software commonly used in neuroimaging. With a single command, FreeSurfer preprocesses an image and outputs several metrics. We analyze gray matter volume, cortical thickness, and surface area since these measures are often used in clinical research (Mitchell et al., 2021). Based on the notion that software variability impacts the estimation of structural measures (Bhagwat et al., 2021; Gronenschild et al., 2012) we hypothesized that there would be differences between FreeSurfer versions in the estimation of cortical thickness, volume, and surface area in a sample of patients with PD and healthy older adults. Second, we investigated whether structural MRI measurement differences between software versions would impact the relationships between brain structure and standard clinical outcomes measured in patients with Parkinson’s disease. Clinical research questions were derived from previous studies on MRI-derived biomarkers of Parkinson’s disease (Hanganu et al., 2014; Mak et al., 2015; for review see Mitchell et al., 2021). More specifically, Hanganu et al. (2014) investigated the group differences between healthy controls, PD with mild cognitive impairment (MCI), and PD without MCI in the rate of change of cortical thickness over time. They also tested the longitudinal group differences in the change of subcortical volumes. The relationship between these structural changes and cognition was also investigated (Hanganu et al., 2014). Similarly, Mak et al. (2015) studied the baseline and longitudinal group differences in cortical thickness and subcortical volumes. The goal of our study is not to reproduce the aforementioned studies but to use previous research as case studies to investigate software variability. We hypothesized that the software version would impact the group differences between healthy controls and PD patients in subcortical volumes and cortical thickness (at baseline and longitudinally). We also hypothesized that software variability would impact the correlation between disease severity and subcortical volumes and cortical thickness in patients with PD (at baseline and longitudinally).

## 2. Methods

### 2.1. Participants

210 patients with PD (181 PD-non-MCI and 29 PD-MCI) and 107 healthy controls (HC) took part in the study. All participants were selected from the Parkinson’s Progression Markers Initiative (PPMI; www.ppmi-info.org). Inclusion criteria for PD patients included the primary diagnosis of PD, available T1-weighted scan, and lack of other neurological diagnosis. One image from PD-non-MCI and one from HC group failed preprocessing. The final sample was 209 PD patients and 106 HC. Out of all participants, 125 PD-non-MCI patients and 106 HC were selected for the clinical part of the analysis. All subjects in this cohort had two study visits with T1-weighted images available, and patients with PD had Unified Parkinson’s Disease Rating Scale (UPDRS) scores. Patients’ (n = 125; 67.2% male) age range was 39.2 - 83.3 (M_age_ = 61.1; SD = 9.3), mean years of education 16.1 ± 3; mean time difference between the visits 1.4 ± 0.7 years; mean UPDRS 23.8 ± 10.2 at baseline and 26.3 ± 11.6 at the follow up visit. Patients with PD-MCI were excluded from the clinical analyses due to the potential confound of MCI on clinical measures such as disease severity. Additional cohort of 160 PD patients (57 women and 103 men; M_age_ = 64.69; SD_age_ = 9.26) has been selected from the Quebec Parkinson Network (QPN) dataset. The inclusion criteria were the diagnosis of PD and availability of structural MRI scan. This sample was used as a replication cohort to test the replicability of the obtained results. Descriptive statistics for all cohorts are reported in Table 1. This study has been conducted in accordance with the Declaration of Helsinki and was exempt from Concordia University’s Research Ethics Unit.

**Table 1.**
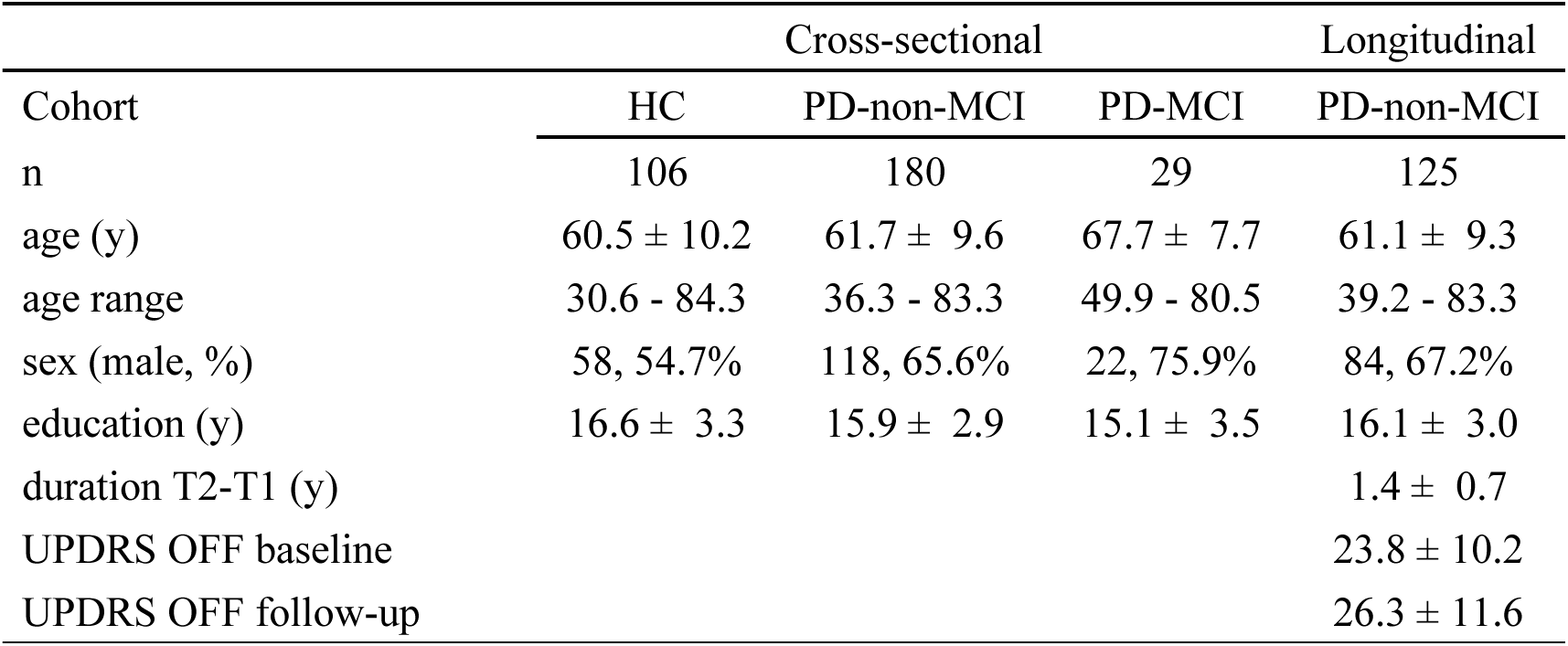
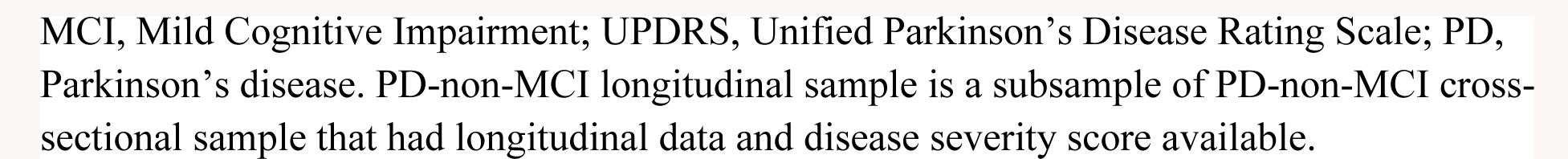
Descriptive statistics.

| Cohort | Cross-sectional |  |  | Longitudinal |
| --- | --- | --- | --- | --- |
|  | HC | PD-non-MCI | PD-MCI | PD-non-MCI |
| n | 106 | 180 | 29 | 125 |
| age (y) | $60.5 \pm 10.2$ | $61.7 \pm 9.6$ | $67.7 \pm 7.7$ | $61.1 \pm 9.3$ |
| age range | 30.6 - 84.3 | 36.3 - 83.3 | 49.9 - 80.5 | 39.2 - 83.3 |
| sex (male, %) | 58, 54.7% | 118, 65.6% | 22, 75.9% | 84, 67.2% |
| education (y) | $16.6 \pm 3.3$ | $15.9 \pm 2.9$ | $15.1 \pm 3.5$ | $16.1 \pm 3.0$ |
| duration T2-T1 (y) | | | | $1.4 \pm 0.7$ |
| UPDRS OFF baseline | | | | $23.8 \pm 10.2$ |
| UPDRS OFF follow-up | | | | $26.3 \pm 11.6$ |

MCI, Mild Cognitive Impairment; UPDRS, Unified Parkinson's Disease Rating Scale; PD, Parkinson's disease. PD-non-MCI longitudinal sample is a subsample of PD-non-MCI cross-sectional sample that had longitudinal data and disease severity score available.

### 2.2. Image acquisition and preprocessing

T1-weighted MRI images were obtained from PPMI that uses standardized acquisition parameters: repetition time = 2300 ms, echo time = 2.98 ms, inversion time = 0.9 s, slice thickness = 1 mm, number of slices = 192, field of view = 256 mm, and matrix size = 256 × 256. However, since PPMI is a multisite project, there may be slight differences in the sites’ setup. T1-weighted MRI images were taken from QPN as a replication data with the following parameters: repetition time = 2300 ms, echo time = 2.98 ms, slice thickness = 1 mm, number of slices = 131, matrix size = 256 x 256.

Brain images were processed using FreeSurfer 5.3, 6.0.1, and 7.3.2 (referred to as FS5, FS6, FS7, accordingly). FreeSurfer’s recon-all function was used for cortical reconstruction. Volumes, cortical thickness, and surface area were extracted for each participant. The longitudinal preprocessing stream was used to analyze images in the analyses that involved clinical data (Reuter et al., 2012). The two timepoints were processed cross-sectionally with the default pipeline, an unbiased template from the two images was created, and data were processed longitudinally. Specifically, an unbiased within-subject template space and image (Reuter & Fischl, 2011) is created using robust, inverse consistent registration (Reuter et al., 2010). Analyses were performed on the Narval cluster hosted at Calcul Québec and part of Digital Research Alliance of Canada. Quality control of raw images was performed with MRIQC. Preprocessed images were visually inspected for quality by assessing brain parcellation and segmentation.

### 2.3. Statistical analyses

#### Stage I: structural MRI measurement differences between software versions

Statistical analyses were divided into two stages. In the first stage, we investigated the differences between FreeSurfer versions in estimating brain volumes, cortical thickness, and surface area in the PPMI cohort. Cortical and subcortical volumes as well as cortical surface areas and thickness were extracted from each T1w scan for all FreeSurfer versions using Destrieux 2009 atlas implemented in FreeSurfer. Paired t-test was used to test the differences between software versions (i.e., FS7 vs FS6, FS7 vs FS5, and FS6 vs FS5) across the entire cohort. Additional analyses were run separately for PD and HC groups. To test for differences in software variability between the two groups, a two-sample t-test was used.

Replication analyses were run with the QPN cohort. In line with the previous analyses, paired t-test was used to test the differences between FreeSurfer versions. The group differences between the PPMI PD group and the QPN cohort in software variability were also tested.

#### Stage II: impact of software variability differences on the relationships between structural brain measurements and clinical outcomes

In the second stage of the analysis, volume, cortical thickness, and surface area were extracted for both timepoints in the longitudinal group. At the subcortical level we investigated the correlation between UPDRS score and subcortical volumes at baseline, and the existence of group differences (PD-non-MCI vs HC) in subcortical volumes. Longitudinal analyses were run to test for the correlation between the rate of change of UPDRS score and the rate of change in subcortical volumes, as well as for group differences in the rate of change of subcortical volumes. Rate of change of subcortical volumes was calculated as [(volume 2 - volume 1) / volume 1] * 100. Partial correlation was used in the correlation analyses and ANCOVA was used in the group analyses. All analyses were controlled for age and sex, and longitudinal analyses were additionally controlled for the time difference between the two visits.

At the vertex level, we examined the correlation between UPDRS scores and baseline cortical thickness, as well as any group differences in cortical thickness at baseline. We conducted additional longitudinal analyses to test the vertex-wise correlation between changes in UPDRS scores and the rate of change in cortical thickness, as well as any vertex-wise group differences in the rate of change in cortical thickness. Rate of change of the cortical thickness in the vertex-wise analyses was calculated as (thickness 2- thickness 1) / (time 2 - time 1), that is mm/year. Age and sex were added as covariates in the vertex-wise analyses. Longitudinal analyses were additionally controlled for the time between the two visits. Cluster-wise *p*-value threshold was used at the *p* < .05 level.

### 2.4. Code availability

We used publicly available software to conduct our study. Pandas v.1.5.2 was used to define cohorts from PPMI data files. FreeSurfer 5.3, 6.0.1, and 7.3.2 were used for image preprocessing and vertex-wise analyses. We used containerized versions of FreeSurfer managed by Boutiques v.0.5.25 (doi:10.5281/zenodo.3839009). The containerized FreeSurfer analyses were executed through the Slurm batch manager on the Narval cluster (https://docs.alliancecan.ca/wiki/Narval/en) hosted at Calcul Québec and part of Digital Research Alliance of Canada. Pearson correlation and t-test analyses were run using SciPy v.1.9.3. Partial correlation and ANCOVA analyses were run using Pingouin 0.5.3. Sørensen–Dice coefficients were calculated using FreeSurfer’s mri_seg_overlap function. Quality control was performed with MRIQC 22.0.1. The code and results are publicly available at https://github.com/LivingPark-MRI/freesurfer-variability with a notebook detailing the analyses. Data used in the notebook were downloaded directly from the PPMI and cannot be shared publicly due to its Data Usage Agreements preventing republishing data. We developed a Python package (LivingPark utils, available at https://github.com/LivingPark-MRI/livingpark-utils) to download and manipulate PPMI data directly from the original PPMI database. As a result, our notebook can be re-executed by anyone with a PPMI account.

## 3. Results

PD patients and HC did not differ in age (*p* > .05) but there were significant differences in years of education (*t* = 2.04; *p* = .04) and the sex frequency (chi2 = 4.02; *p* = .04). Clinical cohort of PD-non-MCI patients and HC group did not differ in age, education, time difference between the two visits, and the sex frequency (*p*s > .05). Descriptive statistics are reported in Table 1.

Quality control of the images indicated that HC and PD groups did not differ in most quality metrics such as coefficient of joint variation, contrast-to-noise ratio, foreground-to-background energy ratio, residual partial gray matter volume effect, Dietrich’s gray matter and total signal-to-noise ratio (*p*s > .05). However, there were significant differences in the gray matter (*p* = .004) and total (*p* = .001) signal-to-noise ratio between the two groups. Quality metrics more often correlated with volume and cortical thickness than surface area. Correlation results for the main sample are reported in Tables S8-S16, for PD in PPMI in Tables S17-S25, and for PD in QPN in Tables S26-34. Images from PD patients in the replication sample differed from PD patients in the main sample in all of the aforementioned quality metrics (*p* < .05). PD patients in QPN, compared to PPMI, had lower coefficient of joint variation and Mortamet’s quality index 1, but higher contrast-to-noise ratio, residual partial gray matter volume effect, gray matter and total signal-to-noise ratio, Dietrich’s gray matter and total signal-to-noise ratio, and Mortamet’s quality index 2.

### 3.1. Stage I: structural MRI measurement differences between software versions

We tested differences between FS versions in the estimation of volume, surface area, and cortical thickness across the sample. Results, reported in Tables 2 and 3, indicate high software variability in the majority of the regions. Statistics are reported in Table S1. Results are reported separately for PD and HC groups in Table S2 and S3.

**Table 2.**
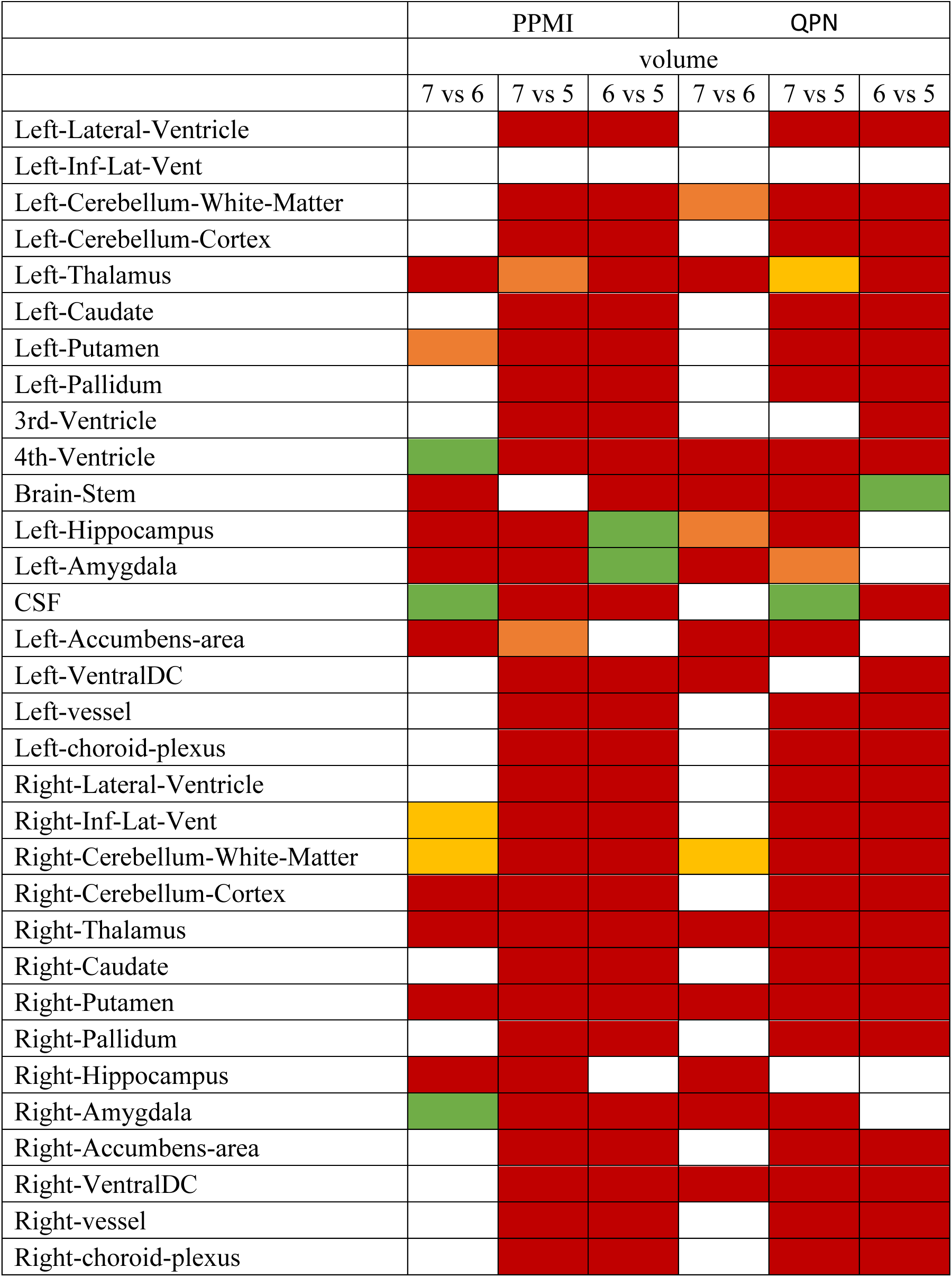
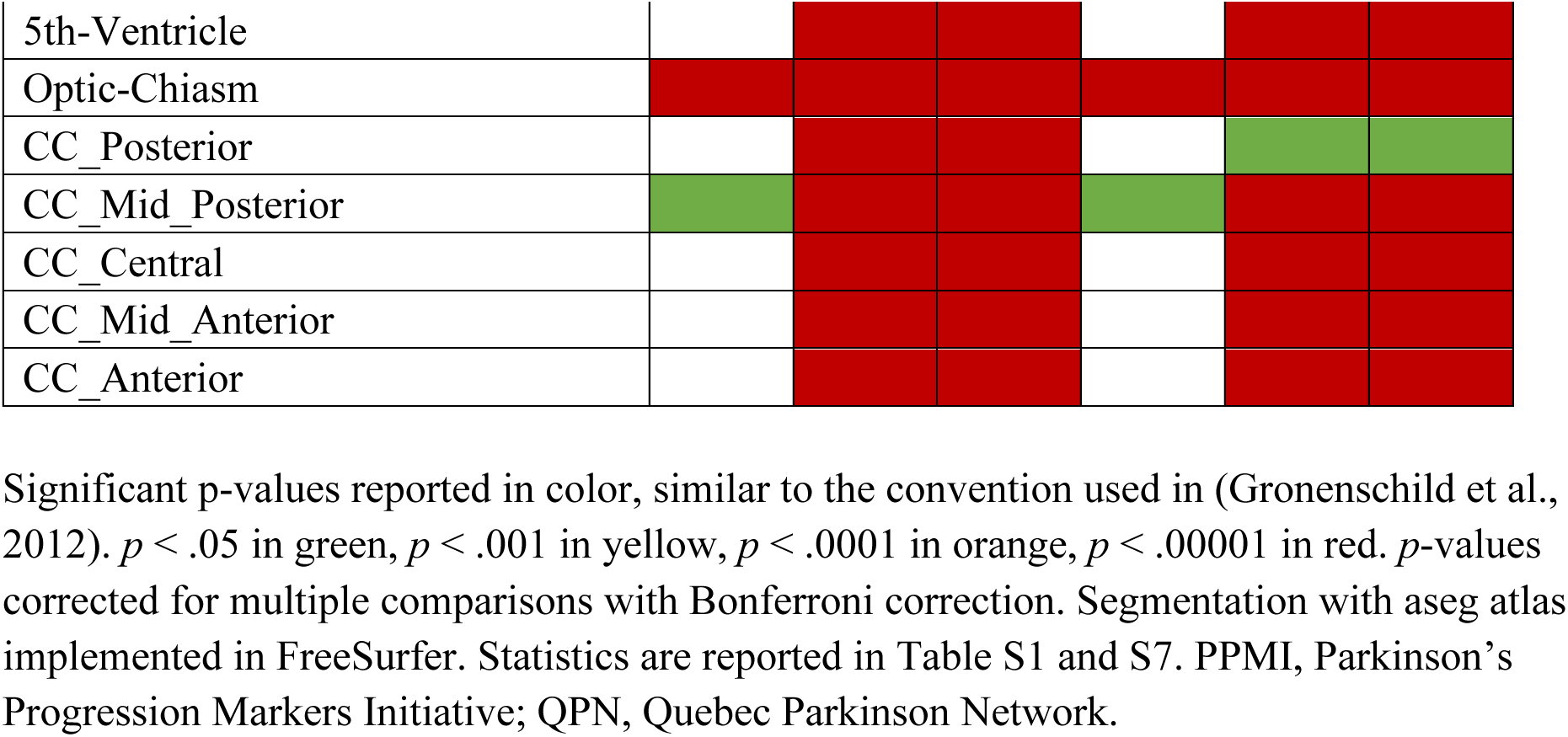
Differences between FreeSurfer versions in estimation of subcortical volume across the PPMI and QPN samples.

**Table 3.**
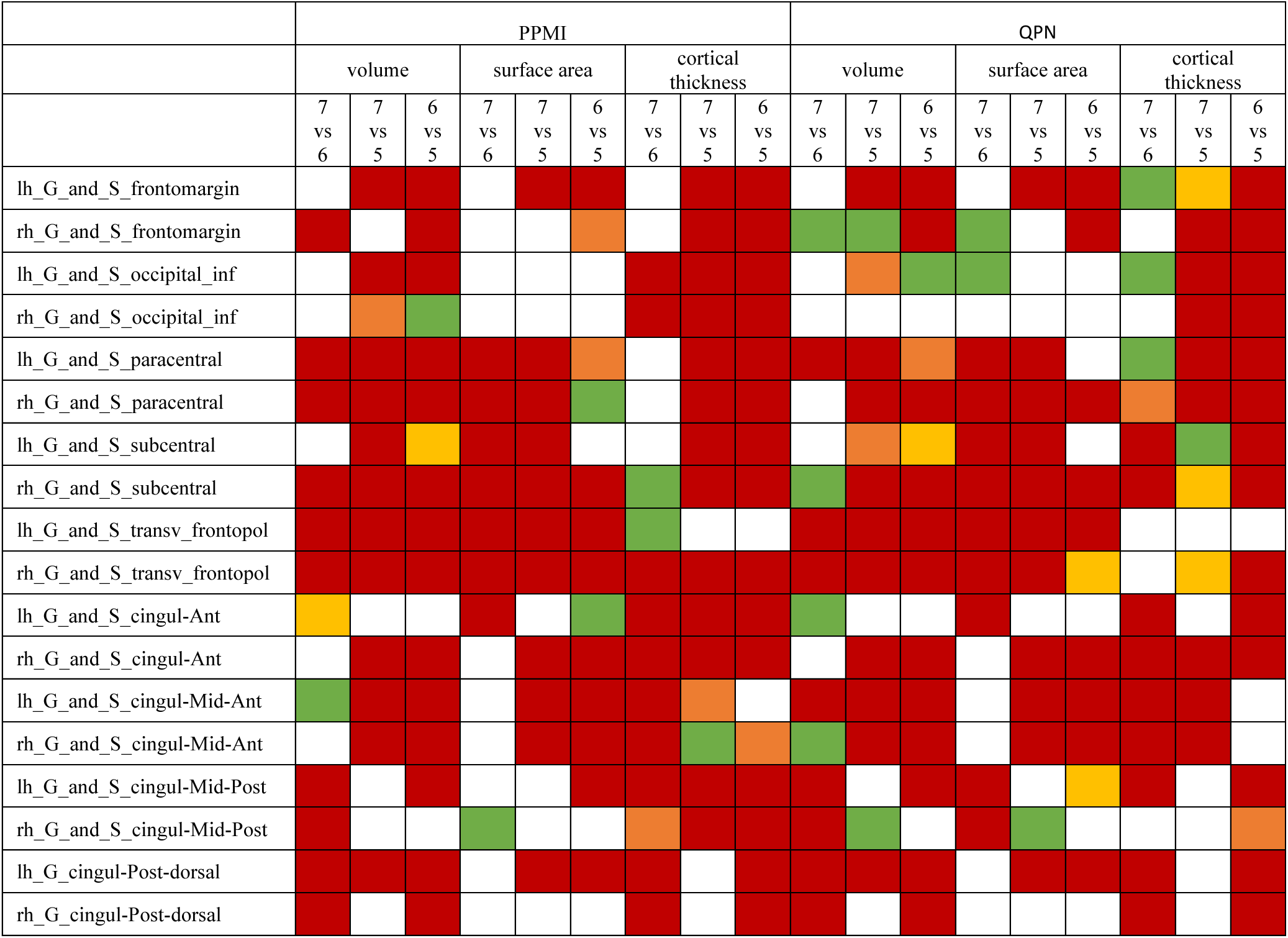
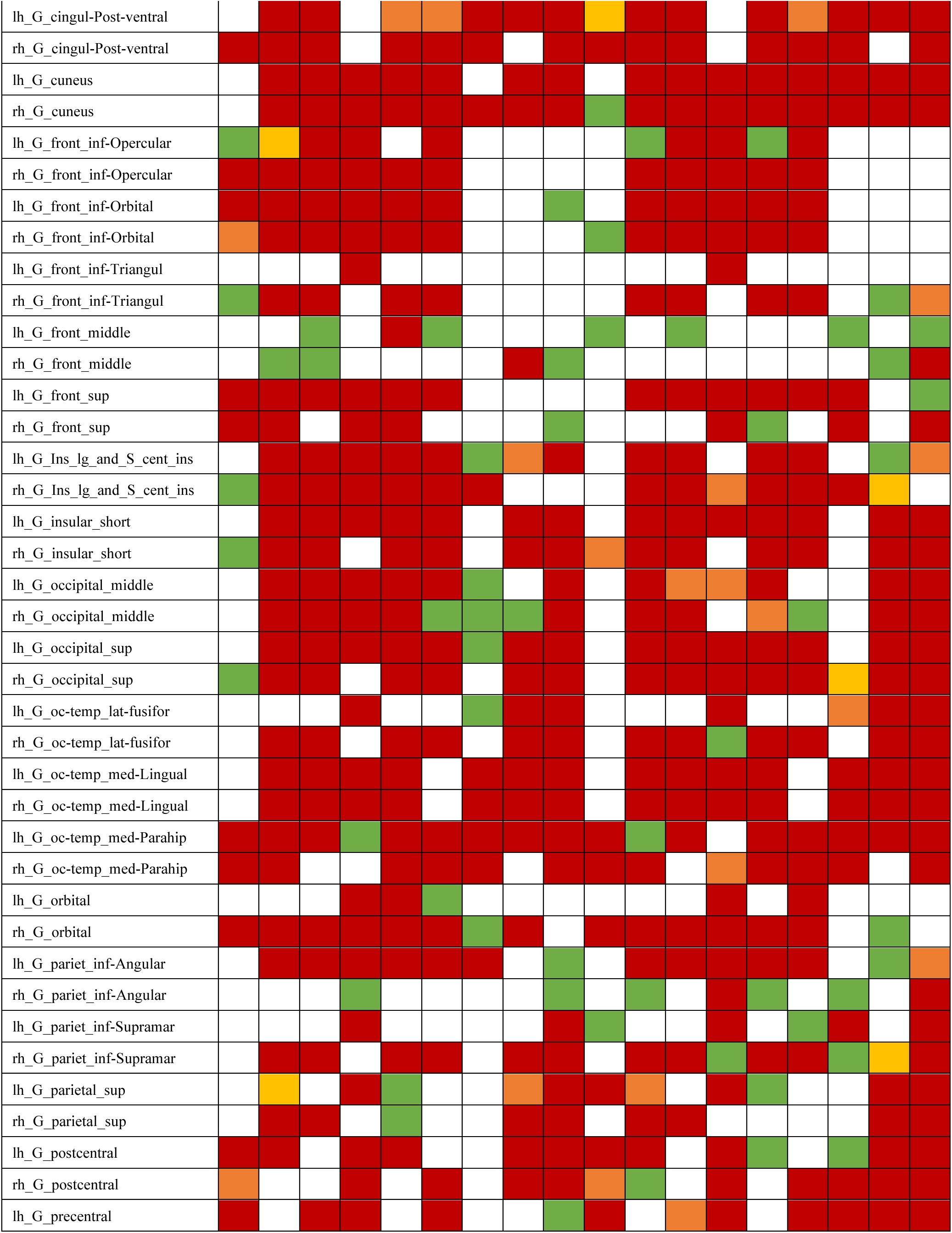
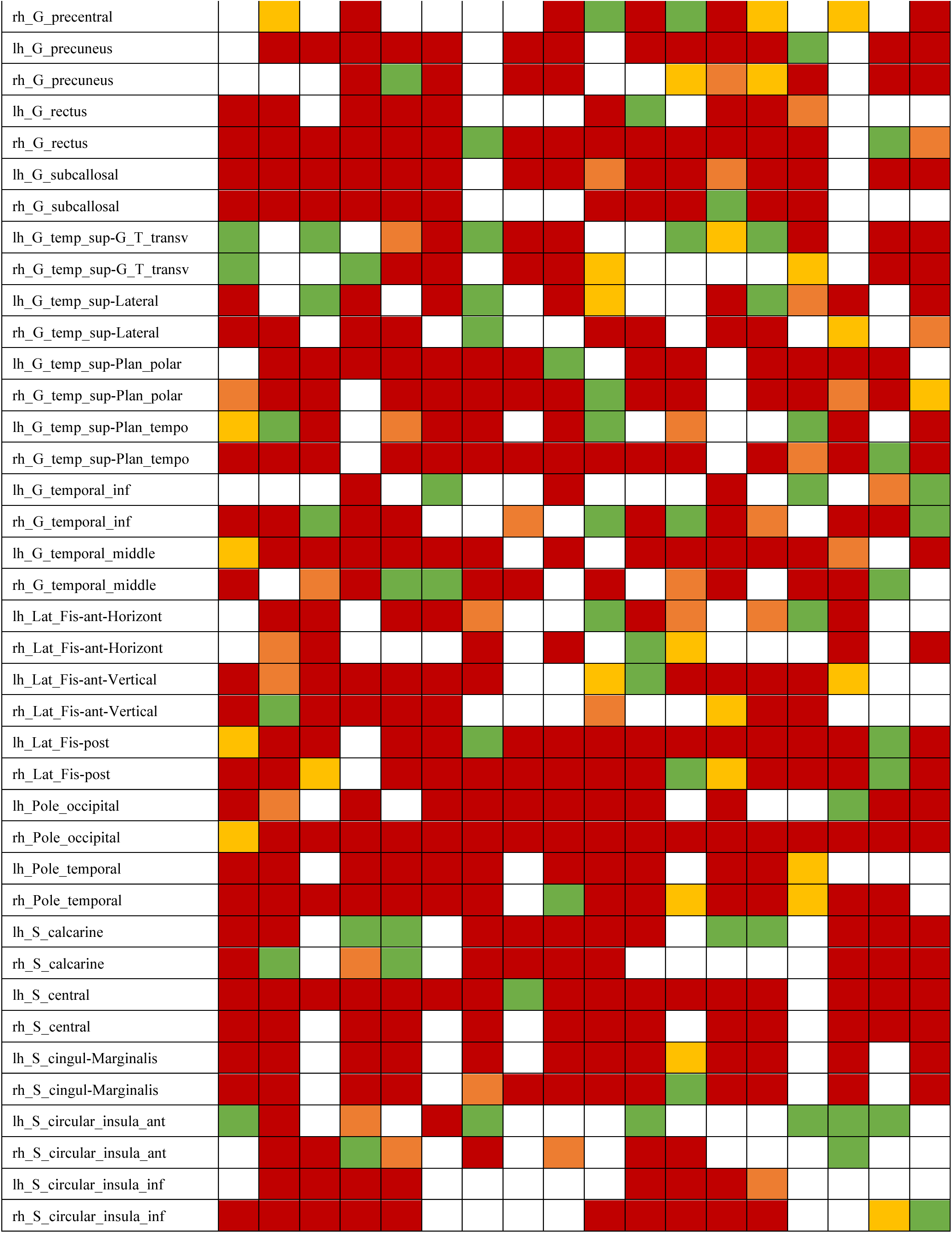
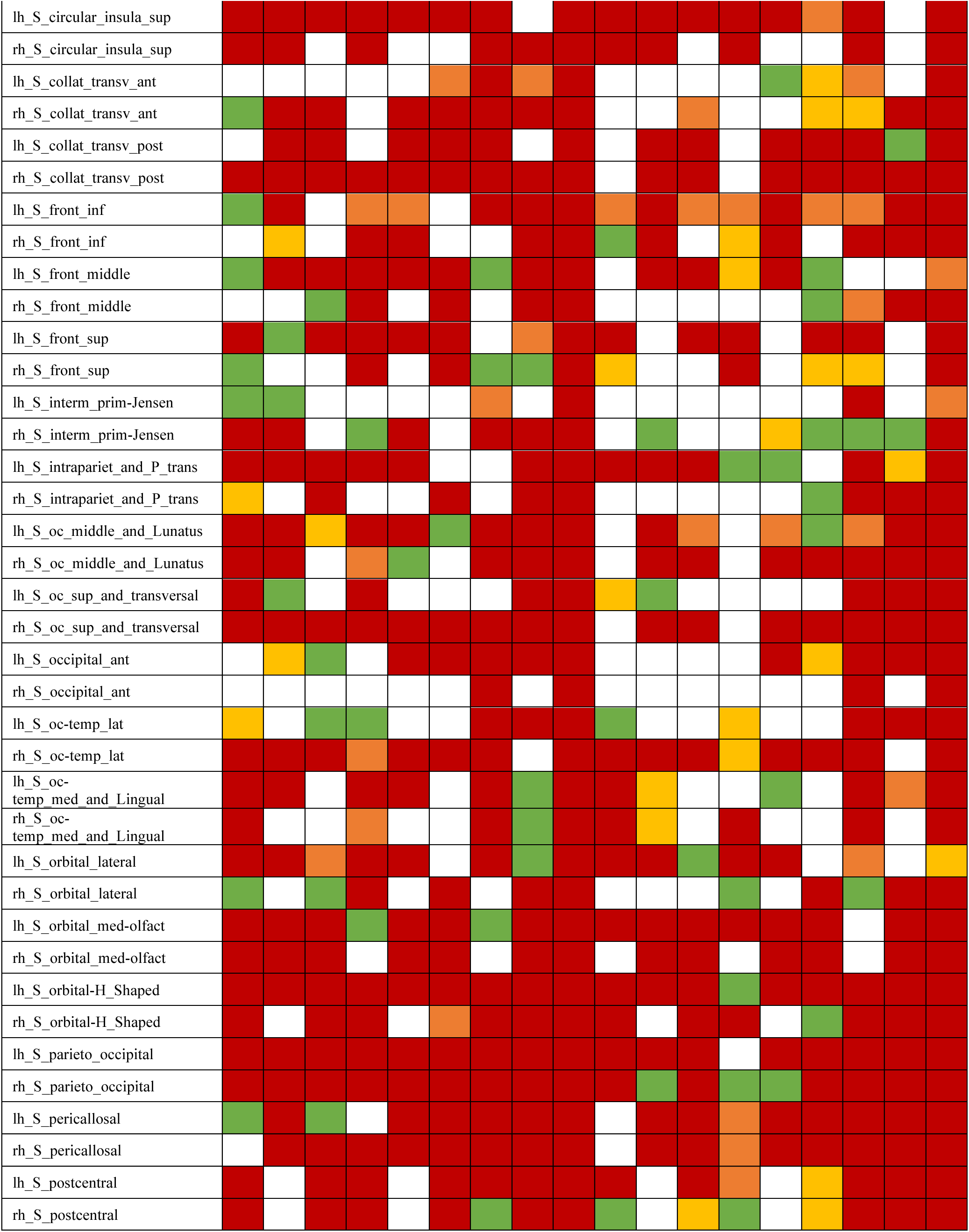
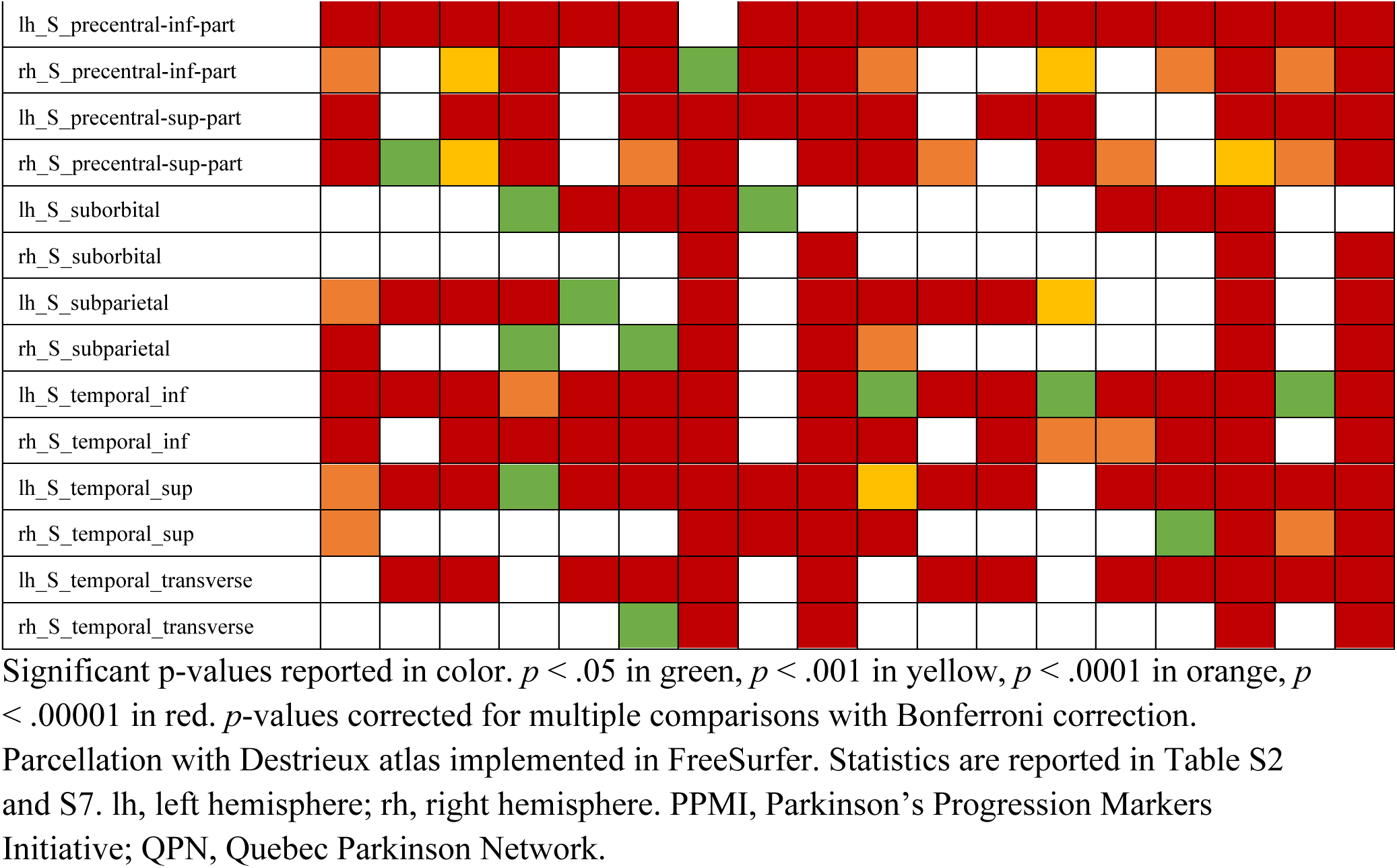
Differences between FreeSurfer versions in estimation of volume, surface area, and cortical thickness across the PPMI and QPN sample.

There were significant differences between FS versions in the estimation of structural metrics based on FreeSurfer’s Destrieux 2009 atlas. FS7 and FS6 differed in the estimation of volume in 62% of the regions, surface in 76%, thickness in 74%; FS7 and FS5 differed in volume in 79%, surface in 73%, thickness in 68%; FS6 and FS5 differed in volume in 75%, surface in 72%, thickness in 86%.

Estimation of volumes, surface areas, and cortical thickness across the three FreeSurfer versions are represented in Figures S1-S4. Between and within-subject variability is represented in Figures 1-4. For certain regions, the differences in the estimation of structural measures are higher for the same subject when analyzed by two different FS versions than for two different subjects analyzed by the same FS version. For example, the left hippocampus had higher between-version variability than between-subject variability (see Figure 1). These variabilities were visualized in Figure S5. The shapes and overlap between hippocampi in two different subjects seem to differ more than in the same subject varying in the software version. This suggests that the volume itself may not be the best metric to measure software variability.

**Figure 1.**
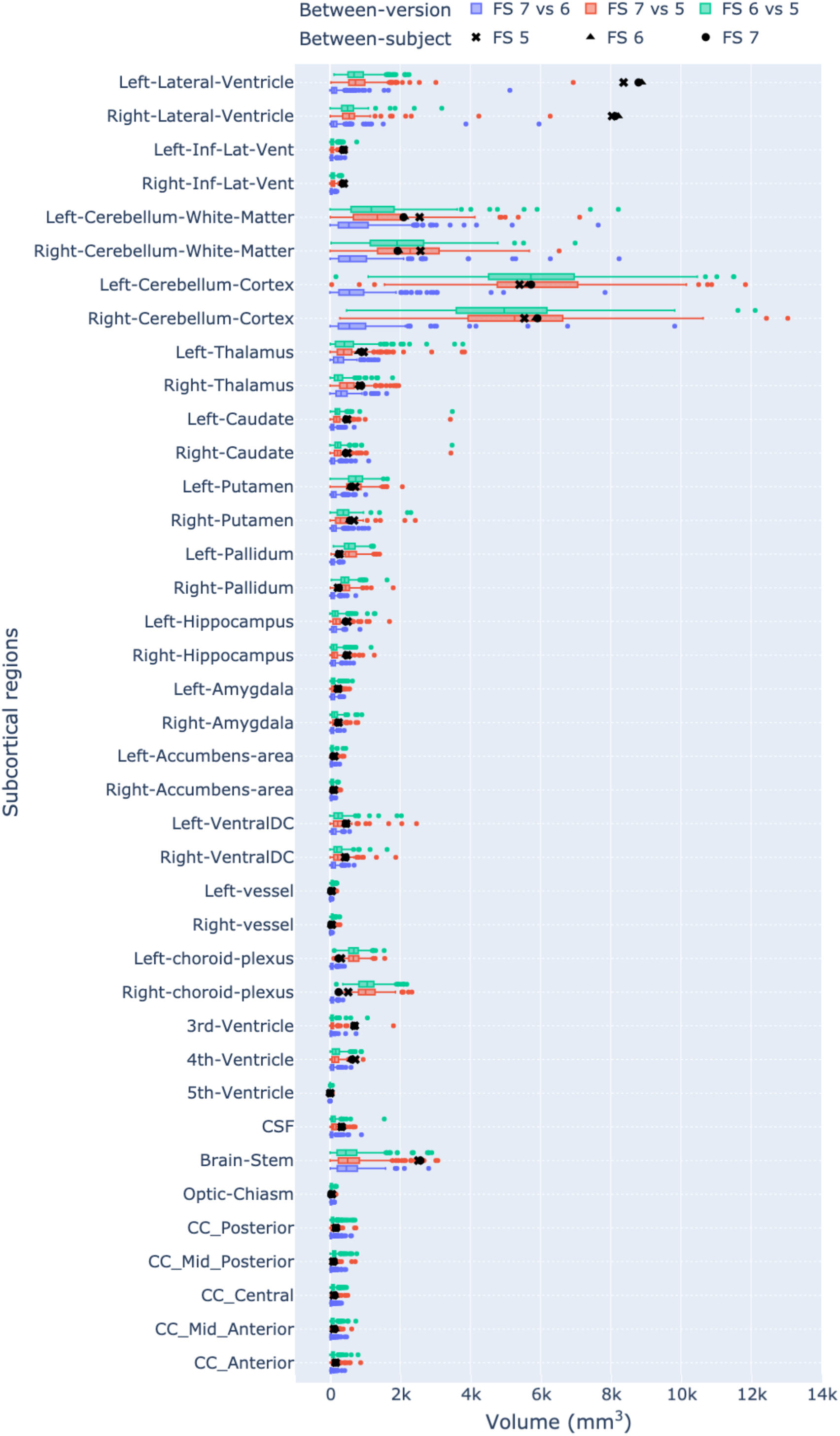
Between-versions (n=3) and between-subject (n=315) subcortical volume variability. Absolute difference between FreeSurfer versions represented as colored box plots, standard deviations represented as black markers. Two outliers not shown in the graph: left-lateral-ventricle (FS 7 vs 5: 31.8k; FS 6 vs 5: 32.4k) and right-lateral-ventricle (FS 7 vs 5: 25.6k; FS 6 vs 5: 26k).

**Figure 2.**
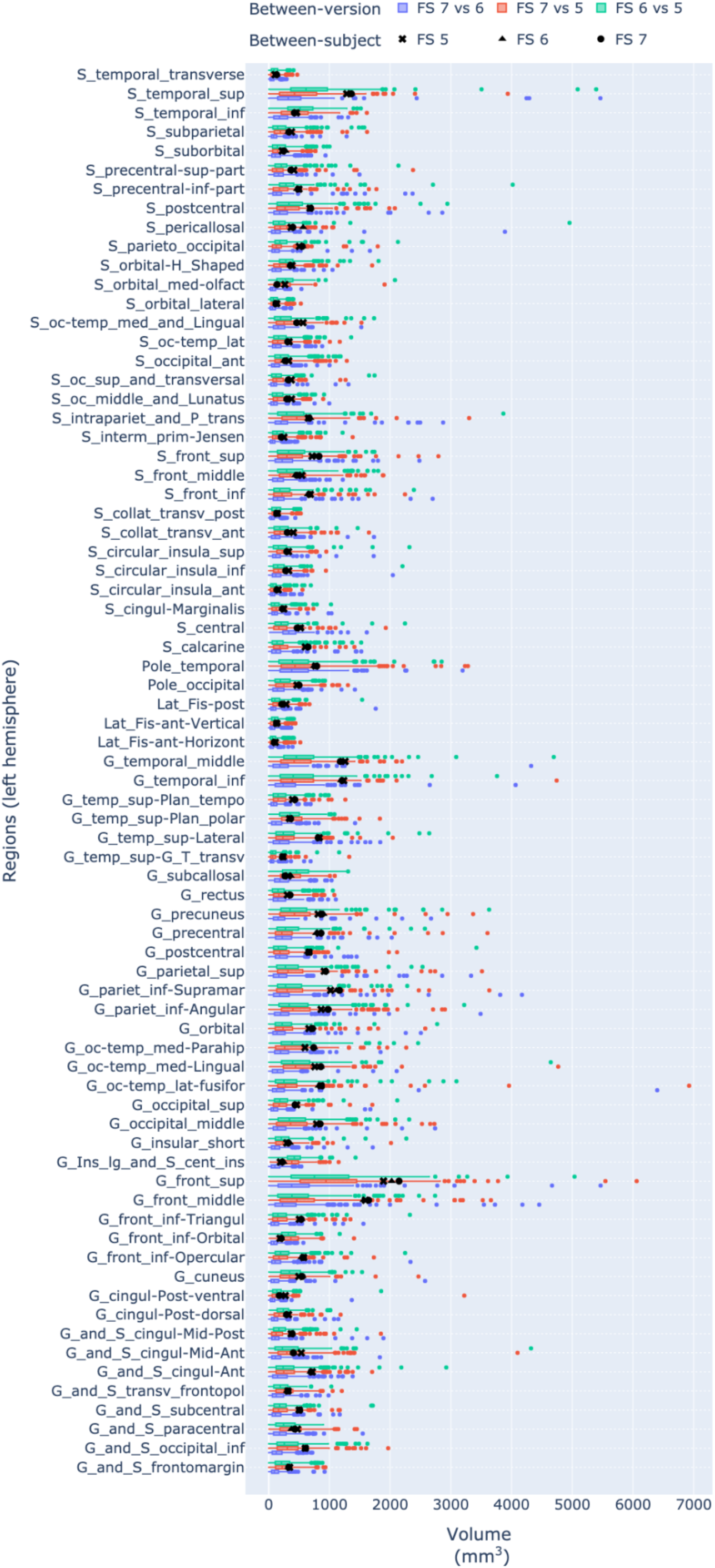
Between-version (n=3) and between-subject (n=315) cortical volume variability in the left hemisphere. Absolute difference between FreeSurfer versions represented as box plots, standard deviations represented as markers. Standard deviation between FreeSurfer versions represented in blue. Results in the right hemisphere reported in Figure S6.

**Figure 3.**
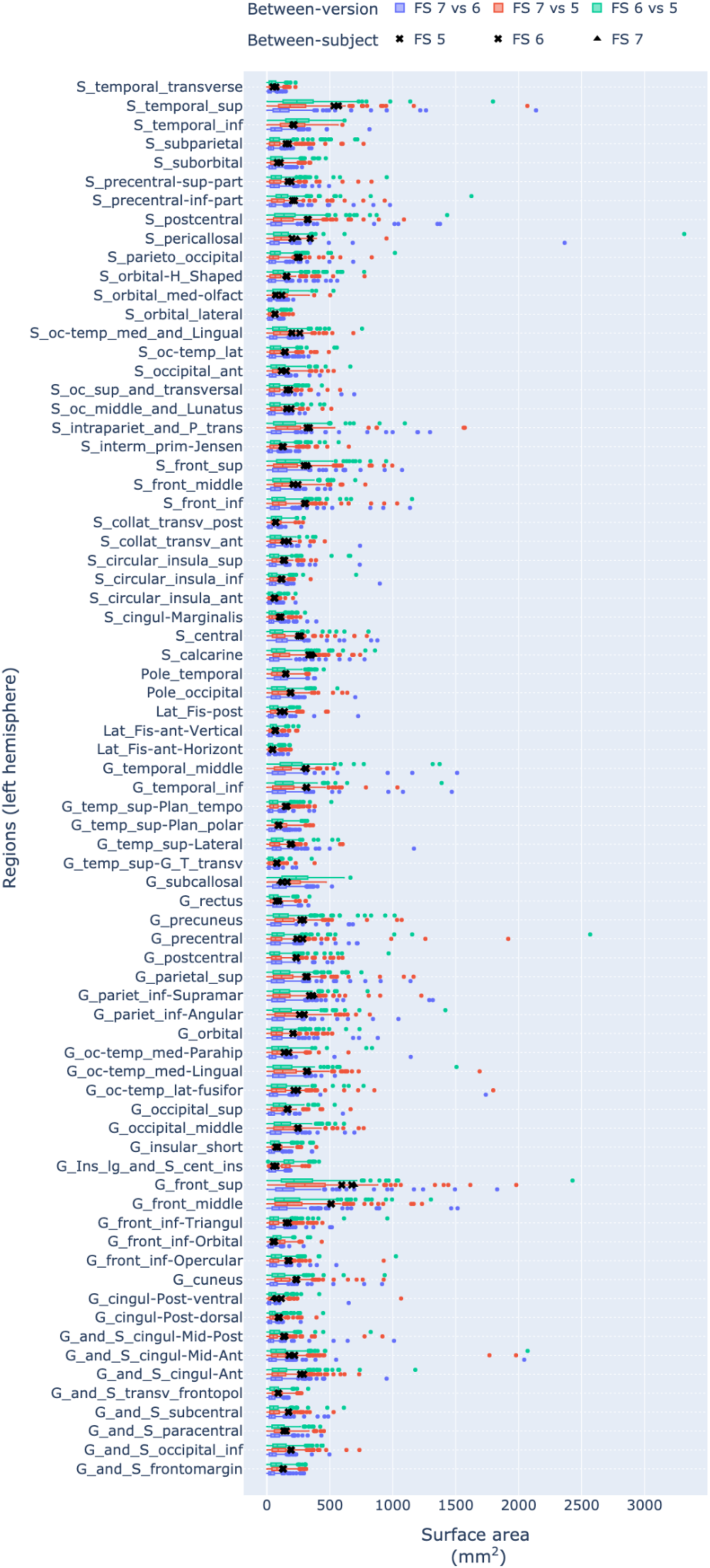
Between-version (n=3) and between-subject (n=315) surface area variability in the left hemisphere. Absolute difference between FreeSurfer versions represented as box plots, standard deviations represented as markers. Standard deviation between FreeSurfer versions represented in blue. Results in the right hemisphere reported in Figure S7.

**Figure 4.**
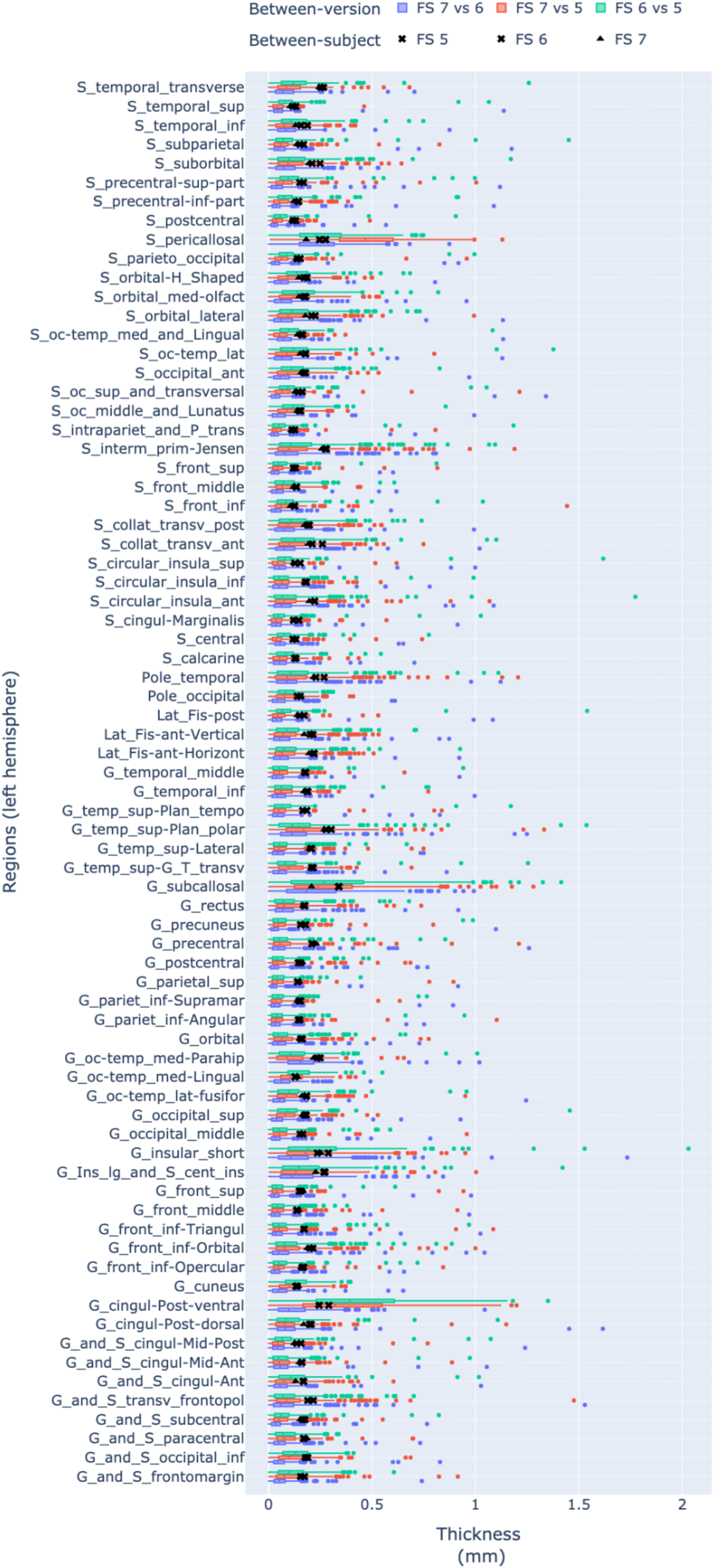
Between-version (n=3) and between-subject (n=315) cortical thickness variability in the left hemisphere. Absolute difference between FreeSurfer versions represented as box plots, standard deviations represented as markers. Results in the right hemisphere reported in Figure S8.

There were no group differences (HC vs PD-non-MCI) in FreeSurfer variability in any of the regions after correction for multiple comparisons (*p* > .05). Results are reported in Table S4.

#### Replication sample

We tested differences between FreeSurfer versions in the estimation of the structural metrics in the replication sample. Results indicate high software variability in the majority of the regions, similarly to the results obtained in the main cohort. Results are reported in Table S7.

There were significant differences between FS versions in the estimation of structural metrics. FS7 and FS6 differed in the estimation of volume in 54% of the regions, surface in 66%, thickness in 73%; FS7 and FS5 differed in volume in 75%, surface in 69%, thickness in 65%; FS6 and FS5 differed in volume in 71%, surface in 67%, thickness in 84%.

### 3.2. Stage II: impact of measurement differences on the relationships between structural brain measurements and clinical outcomes

#### Subcortical analyses

We tested the group differences (PD vs HC) in subcortical volumes at baseline. To quantify the impact of Freesurfer versions on analysis outcomes, we calculated the SD coefficients between (1) the set of regions with significant differences between PD and HC computed with Freesurfer version A, and (2) the set of regions with significant differences between PD and HC computed with Freesurfer version B. For example, the Sørensen–Dice coefficient for the baseline differences between PD and HC in subcortical volumes between FS5 and FS6 was calculated as 2(4)/(4+5) = .89. The Sørensen–Dice coefficient of the significant results between the FS7 and FS6 was .77, between FS7 and FS5 was .67, and between FS6 and FS5 was .89. Results are reported in Table 4.

**Table 4.**
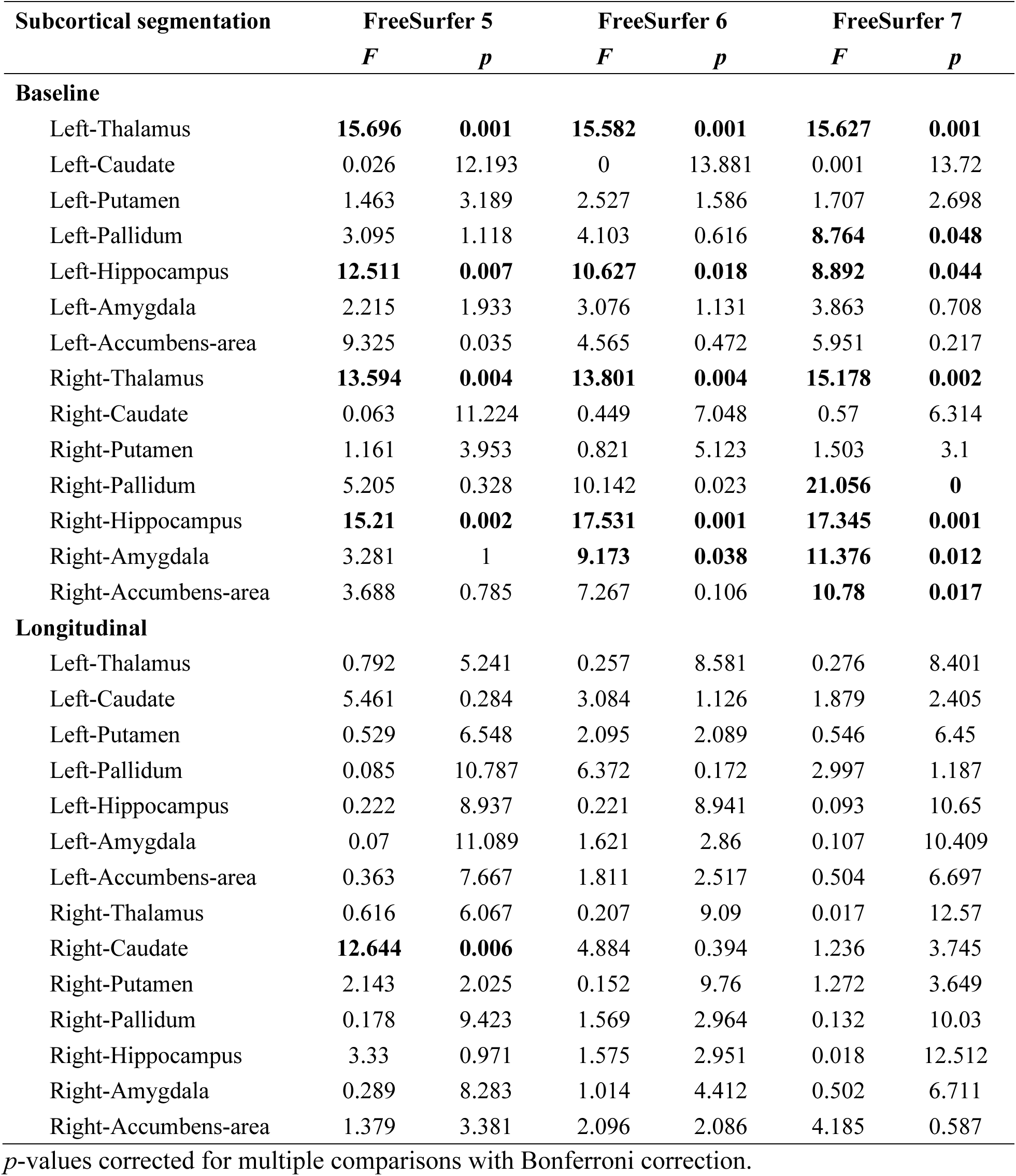
Group differences (PD vs HC) in subcortical volumes at baseline and longitudinal group differences in the rate of change in subcortical volumes.

We investigated group differences (PD vs HC) in the rate of change of subcortical volumes. The Sørensen–Dice coefficients of the significant results across all FS versions were zero. Results are reported in Table 4. Software variability has a substantial impact on both baseline and longitudinal analysis, even though the lack of significance in the longitudinal analysis would deserve additional studies.

We investigated correlation between UPDRS score and subcortical volumes at baseline. There was a significant correlation between the right putamen volume and the UPDRS score reported in FS6 and FS7 (*p*s < .05), but not in FS5. Results are reported in Table 5.

**Table 5.** Correlation between the UPDRS score and subcortical volumes at baseline, and longitudinal correlation between the change in UPDRS score and the rate of change in subcortical volumes.

| Subcortical segmentations | FreeSurfer 5 |  | FreeSurfer 6 |  | FreeSurfer 7 |  |
| --- | --- | --- | --- | --- | --- | --- |
|  | *r* | *p* | *r* | *p* | *r* | *p* |
| **Baseline** |  |  |  |  |  |  |
| Left-Thalamus | -0.013 | 12.36 | -0.026 | 10.838 | -0.016 | 12.017 |
| Left-Caudate | -0.206 | 0.312 | -0.178 | 0.685 | -0.182 | 0.611 |
| Left-Putamen | -0.188 | 0.52 | -0.202 | 0.352 | -0.221 | 0.194 |
| Left-Pallidum | -0.182 | 0.622 | -0.085 | 4.928 | -0.15 | 1.357 |
| Left-Hippocampus | -0.124 | 2.426 | -0.11 | 3.144 | -0.12 | 2.589 |
| Left-Amygdala | -0.099 | 3.846 | -0.155 | 1.206 | -0.176 | 0.722 |
| Left-Accumbens-area | 0.127 | 2.27 | 0.085 | 4.921 | 0.086 | 4.853 |
| Right-Thalamus | -0.013 | 12.416 | -0.033 | 10.04 | -0.058 | 7.311 |
| Right-Caudate | -0.11 | 3.15 | -0.107 | 3.322 | -0.118 | 2.687 |
| Right-Putamen | -0.222 | 0.192 | **-0.269** | **0.037** | **-0.263** | **0.047** |
| Right-Pallidum | -0.015 | 12.149 | 0.041 | 9.142 | -0.007 | 13.181 |
| Right-Hippocampus | -0.223 | 0.183 | -0.163 | 1.014 | -0.189 | 0.515 |
| Right-Amygdala | -0.159 | 1.111 | -0.156 | 1.201 | -0.237 | 0.115 |
| Right-Accumbens-area | 0.001 | 13.847 | 0.025 | 11.028 | -0.034 | 9.929 |
| **Longitudinal** |  |  |  |  |  |  |
| Left-Thalamus | 0.14 | 1.737 | 0.037 | 9.654 | -0.012 | 12.556 |
| Left-Caudate | 0.096 | 4.124 | -0.016 | 11.999 | -0.002 | 13.744 |
| Left-Putamen | -0.007 | 13.159 | 0.1 | 3.812 | -0.011 | 12.63 |
| Left-Pallidum | 0.022 | 11.384 | 0.145 | 1.551 | 0.039 | 9.401 |
| Left-Hippocampus | 0.067 | 6.522 | 0.066 | 6.567 | 0.063 | 6.829 |
| Left-Amygdala | 0.103 | 3.65 | -0.076 | 5.715 | 0.011 | 12.64 |
| Left-Accumbens-area | -0.101 | 3.734 | 0.076 | 5.662 | 0.091 | 4.485 |
| Right-Thalamus | -0.009 | 12.956 | -0.016 | 12.042 | -0.016 | 12.01 |
| Right-Caudate | -0.122 | 2.523 | -0.221 | 0.204 | -0.229 | 0.155 |
| Right-Putamen | 0.062 | 6.985 | 0.208 | 0.305 | 0.141 | 1.714 |
| Right-Pallidum | -0.039 | 9.358 | -0.089 | 4.609 | 0.057 | 7.462 |
| Right-Hippocampus | 0.12 | 2.637 | 0.005 | 13.396 | -0.01 | 12.759 |
| Right-Amygdala | -0.002 | 13.745 | -0.031 | 10.341 | -0.141 | 1.71 |
| Right-Accumbens-area | -0.132 | 2.077 | -0.025 | 10.977 | -0.018 | 11.829 |

*p*-values corrected for multiple comparisons with Bonferroni correction.

We investigated correlation between the change in UPDRS score and the rate of change of subcortical volumes. No result remained significant after Bonferroni correction. Results are reported in Table 5.

#### Vertex-wise analyses

##### Correlation analyses

**Table 6.**
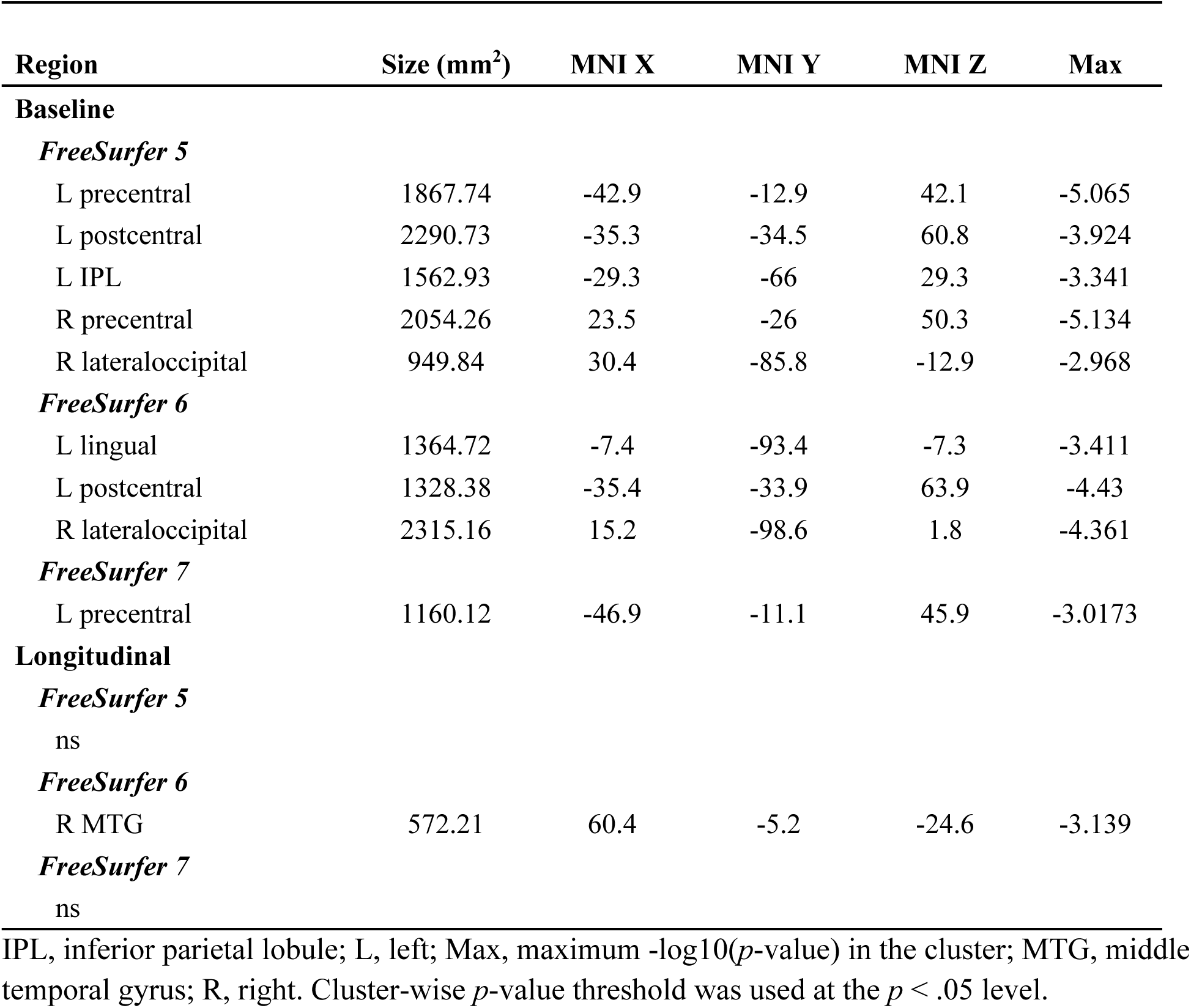
Vertex-wise correlation between UPDRS score and cortical thickness at baseline, and the correlation between the rate of change in UPDRS and the rate of change in cortical thickness.

| Region | Size (mm<sup>2</sup>) | MNI X | MNI Y | MNI Z | Max |
| --- | --- | --- | --- | --- | --- |
| **Baseline** |  |  |  |  |  |
| *FreeSurfer 5* |  |  |  |  |  |
| L precentral | 1867.74 | -42.9 | -12.9 | 42.1 | -5.065 |
| L postcentral | 2290.73 | -35.3 | -34.5 | 60.8 | -3.924 |
| L IPL | 1562.93 | -29.3 | -66 | 29.3 | -3.341 |
| R precentral | 2054.26 | 23.5 | -26 | 50.3 | -5.134 |
| R lateraloccipital | 949.84 | 30.4 | -85.8 | -12.9 | -2.968 |
| *FreeSurfer 6* |  |  |  |  |  |
| L lingual | 1364.72 | -7.4 | -93.4 | -7.3 | -3.411 |
| L postcentral | 1328.38 | -35.4 | -33.9 | 63.9 | -4.43 |
| R lateraloccipital | 2315.16 | 15.2 | -98.6 | 1.8 | -4.361 |
| *FreeSurfer 7* |  |  |  |  |  |
| L precentral | 1160.12 | -46.9 | -11.1 | 45.9 | -3.0173 |
| **Longitudinal** |  |  |  |  |  |
| *FreeSurfer 5* |  |  |  |  |  |
| ns |  |  |  |  |  |
| *FreeSurfer 6* |  |  |  |  |  |
| R MTG | 572.21 | 60.4 | -5.2 | -24.6 | -3.139 |
| *FreeSurfer 7* |  |  |  |  |  |
| ns |  |  |  |  |  |

IPL, inferior parietal lobule; L, left; Max, maximum $-\log_{10}(p\text{-value})$ in the cluster; MTG, middle temporal gyrus; R, right. Cluster-wise $p$ -value threshold was used at the $p < .05$ level.

**Figure 5.**
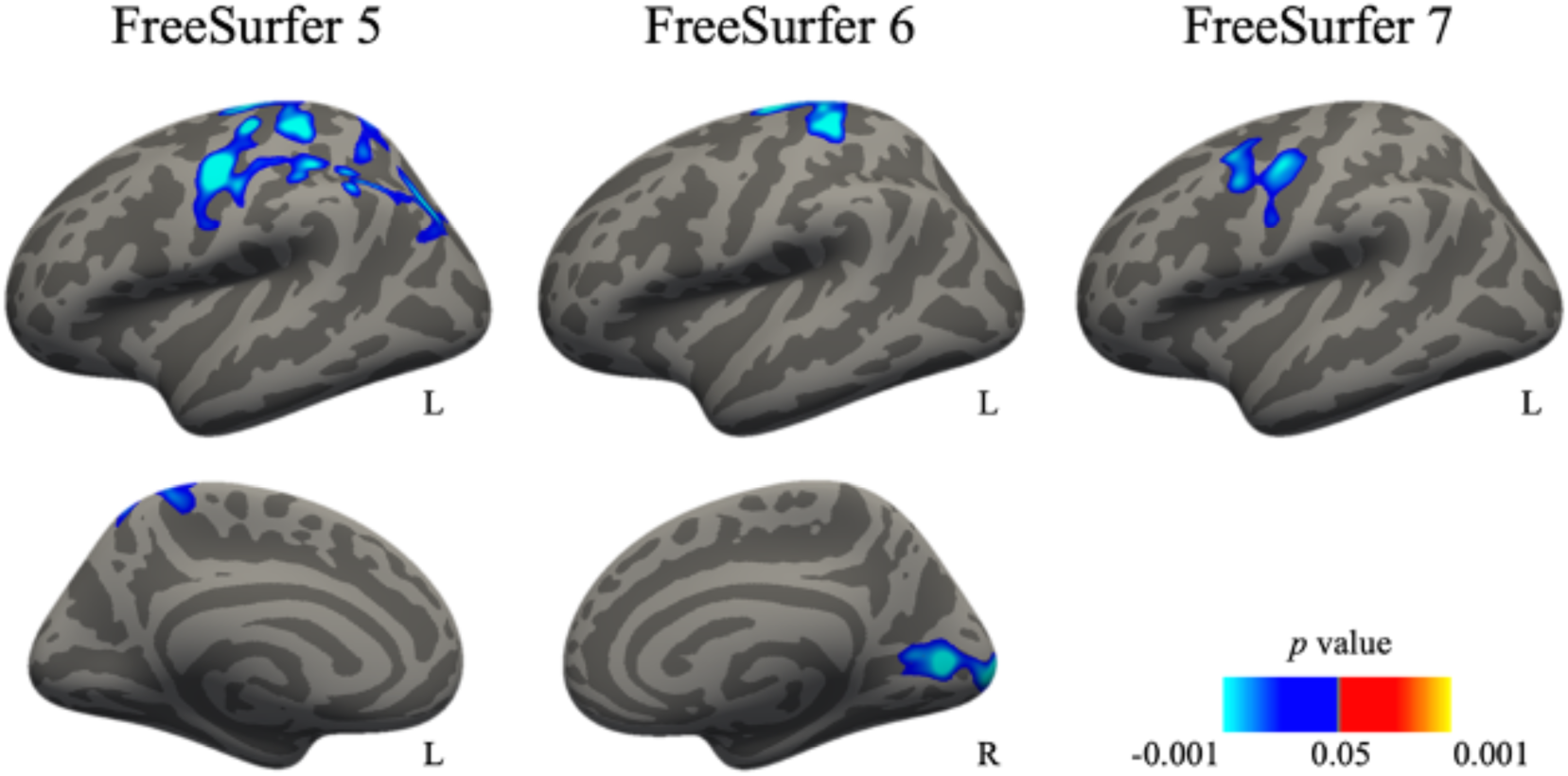
Vertex-wise correlation between UPDRS score and cortical thickness at baseline.

**Figure 6.**
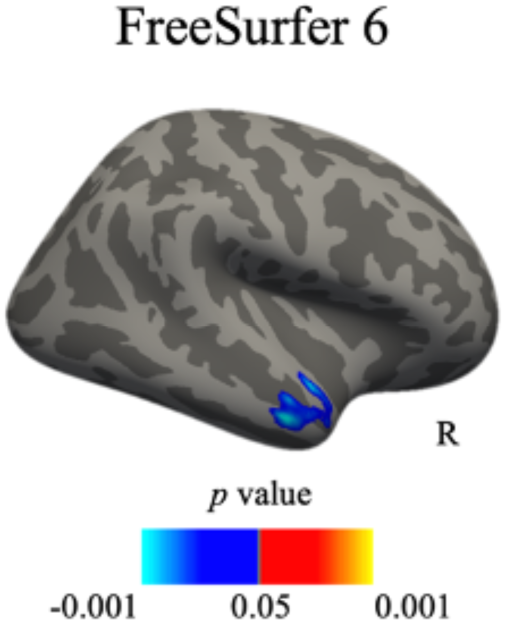
Vertex-wise correlation between the rate of change in UPDRS and the rate of change in cortical thickness. Correlations in FreeSurfer 5 and 7 were not significant.

##### Group analyses

**Table 7.**
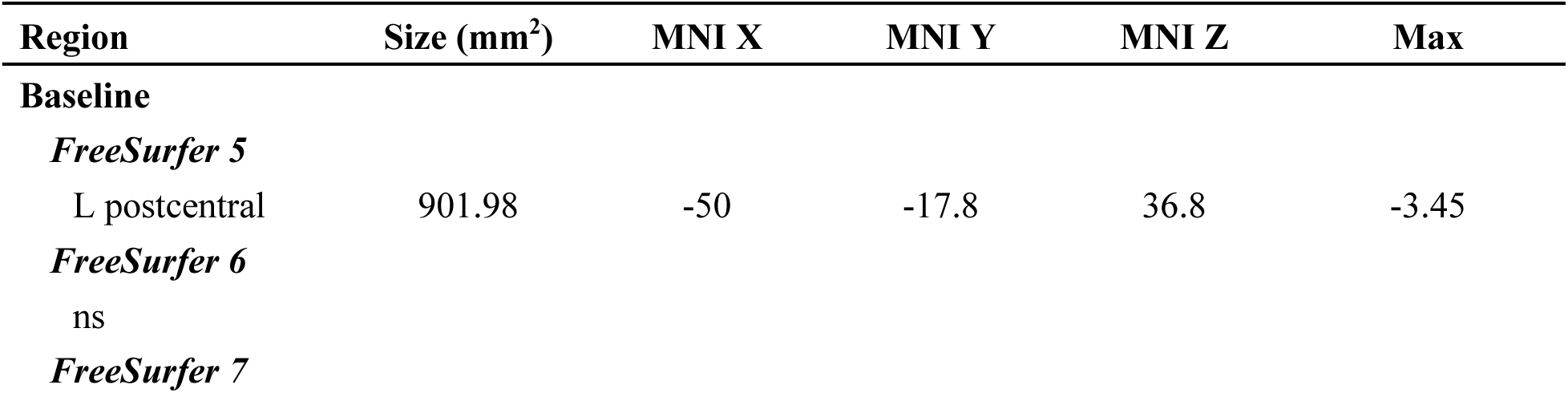
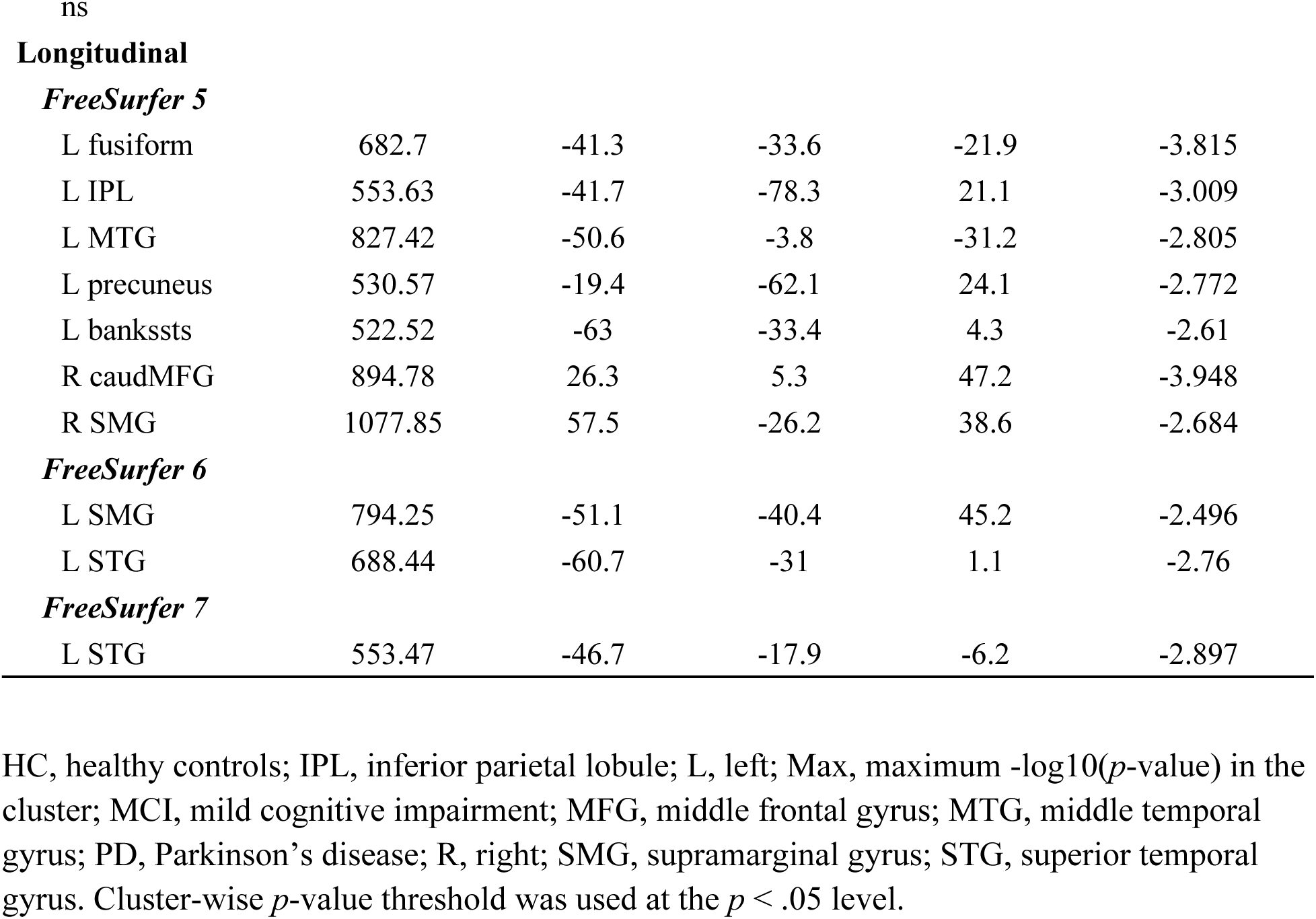
Group differences (HC vs PD-non-MCI) in cortical thickness at baseline and in the rate of change in cortical thickness.

**Figure 7.**
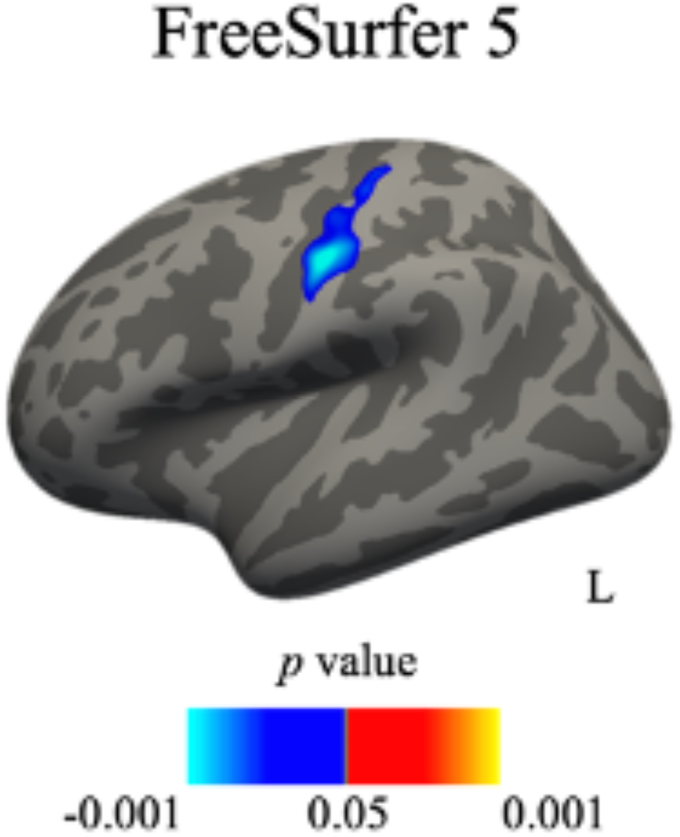
Group differences (HC vs PD-non-MCI) in cortical thickness at baseline. Differences in FreeSurfer 6 and 7 were not significant.

**Figure 8.**
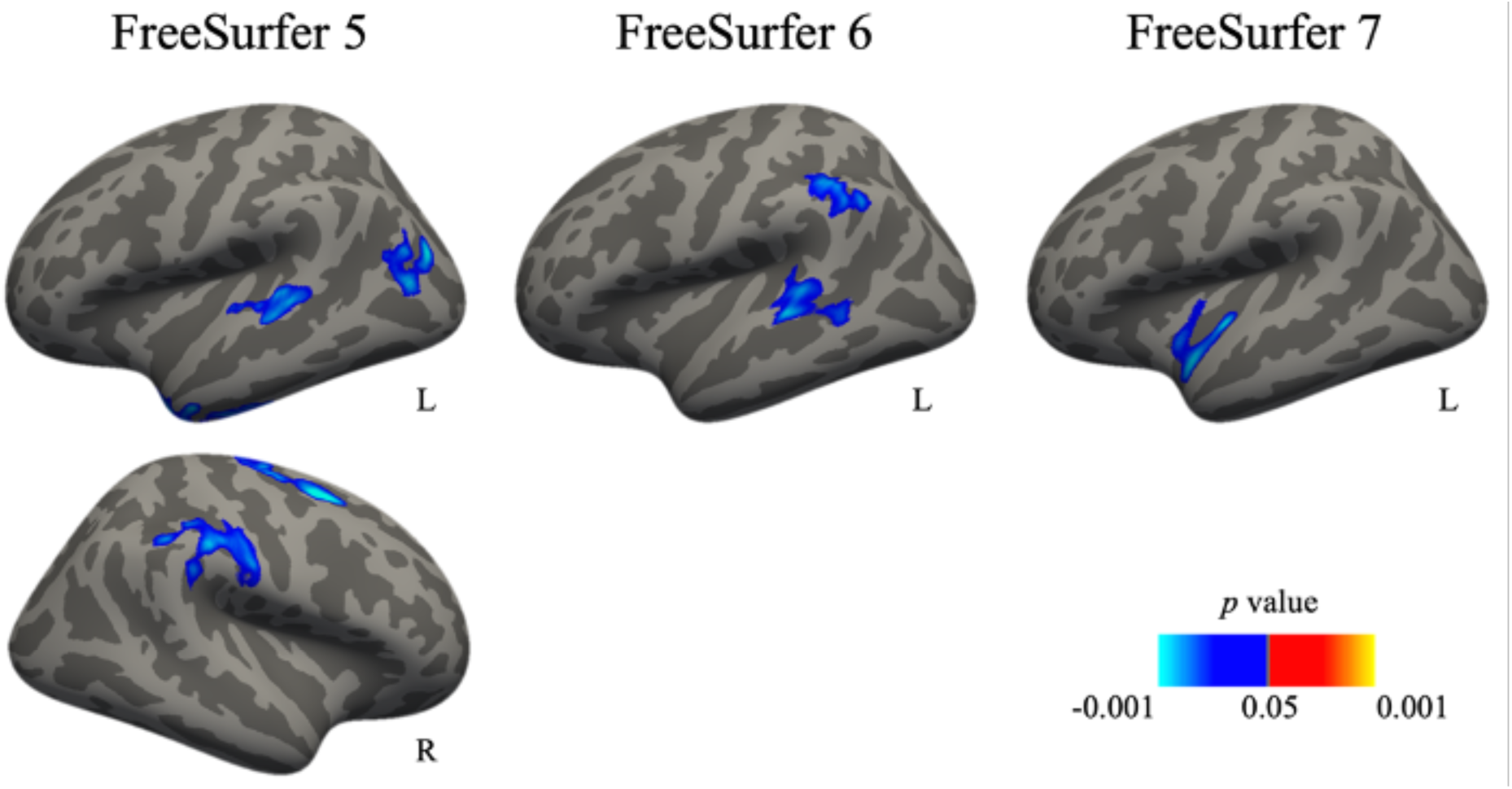
Group differences (HC vs PD-non-MCI) in the rate of change in cortical thickness.

## 4. Discussion

This study investigated the impact of software variability resulting from FreeSurfer version updates on the estimation of MRI-derived structural brain measures and the impact of this measurement variability on the relationships between brain measures and clinical outcomes in Parkinson’s disease. We observed variability across many brain regions in the estimation of volume, surface area, and cortical thickness. The differences were more wide-spread when comparing FS5 to the newer FreeSurfer versions while the differences between FS7 and FS6 were less prevalent, although were still common. Our results are in line with previous research on FreeSurfer software variability (Bhagwat et al., 2021; Gronenschild et al., 2012; Haddad et al., 2023) although these studies focused on different FreeSurfer release versions. Relatively higher similarity between FS7 and FS6 in our study is in line with results reported by Haddad et al. (2023) who report the lowest compatibility with FS5. Our data support the notion that the choice of the software version may impact the estimation of structural measures. Importantly, not all regions were equally impacted by the software variability and we found no differences in some of the regions. Volume estimation of subcortical structures was relatively not impacted by the variability between FS7 and FS6 which is in line with Haddad et al. (2023). This could suggest that researchers focusing solely on these structures or volume measures should not worry about the choice of the software. However, we refrain from such recommendations since there is no clear pattern of regions that are less susceptible to the variability.

The impact of software variability on the estimation of all three structural measures is the same in patients with PD and HC as we did not find any differences between these two groups. It suggests that the variability introduced by different preprocessing toolboxes might also be present in other cohorts. This result supports the fact that existing software variability results (such as Botvinik-Nezer et al., 2020; Filip et al., 2022; Gronenschild et al., 2012) could also apply to the PD populations similar to the PPMI and QPN samples tested in our study.

This may be explained by the fact that our PD sample, compared to HC, did not exhibit significant differences in image quality metrics except for the signal-to-noise ratio. In populations where data quality is reduced compared to HC (e.g., children) software variability might still manifest differently. Nevertheless, data obtained independently in another recent study suggest that software variability does not relate to image quality altogether (Sanz-Robinson et al., 2023). This study found that data quality did not correlate with the difference in the volume estimation of the hippocampus between FreeSurfer and FSL. Importantly, there was a limited correlation between the software variability and quality metrics. Variability correlated in some regions with coefficient of joint variation and contrast-to-noise ratio even though PD and HC groups did not differ in these metrics. On the other hand, there was a correlation between gray matter signal-to-noise ratio in some regions and the two groups differed in this metric. This suggests that the impact of quality metrics on software variability is independent from group differences in quality metrics.

To test the impact of software variability in a clinical setting, we investigated the differences between FreeSurfer versions in the testing of clinical hypotheses. We tested group differences (patients vs healthy participants) in subcortical volumes and cortical thickness as well as the correlation between disease severity in patients and their cortical thickness. These hypotheses were derived from prior neuroimaging studies on PD (Hanganu et al., 2014; Mak et al., 2015; for review see Mitchell et al., 2021). To our knowledge this is the first attempt to assess the degree of impact of software variability on clinical findings in the PD population.

The significance of the differences between PD and HC in the estimation of the subcortical volumes was impacted by software variability. The Sørensen–Dice coefficient of subcortical volumes between FS versions ranged from .67 to .89. Inconsistent results are in line with the study by Filip et al. (2022) who reported variability across FS versions in the group differences in volume estimation in patients with diabetes. In the context of the presence of subcortical atrophy in the PD population (Mitchell et al., 2021), our results indicate that various research teams, addressing the same clinical question, may or may not obtain significant results depending on the software they use. We have also found a significant correlation between the baseline volume of the right putamen and the disease severity in two FS versions. This result was not significant in FS5 which is in line with the results of the first part of our study. We have reported stronger differences between FS5 and the other two FS versions in the right putamen volume estimation than between FS6 and FS7. There are prior studies that indicate the link between the putamen volume and disease severity in PD (Mak et al., 2014; Wilson et al., 2019). Vertex-wise analyses yielded different results depending on the FS version. There was a correlation between the disease severity and cortical thickness in numerous regions in FS5, a few in FS6, and only one significant cluster in FS7. This is not surprising given previous reports showing that FS5 is often less stringent than more recent versions (Filip et al., 2022; Haddad et al., 2023).

Only FS5 reported group differences between HC and PD-non-MCI in cortical thickness at baseline. This again would support the notion that FS5 is more permissive. This was also observed in the longitudinal rate of change in cortical thickness where FS5 gave more significant regions than the other two FS versions. However, we also report results that are significant in versions FS6 and FS7 but not FS5. Furthermore, reporting regions’ labels is not sufficient to determine similarity between the output. For example, both FS6 and FS7 reported significant differences in the left superior temporal gyrus. However, only upon reviewing the activation maps it becomes clear that these two regions are distinct (the former being posterior and the latter anterior) with no overlap between them.

There were several changes in FreeSurfer across subsequent releases. A new fsaverage space was introduced in FS6, along with improved accuracy of the cortex labels and better handling of issues with registration. FS7 brought changes in segmentation and some statistical analysis options were deprecated (e.g., QDEC). These are only a few examples of changes that can partially explain obtained differences even though we used the same pipeline in all our analyses.

Our study comes with limitations. We investigated one release of the major FS versions. Testing more versions, including minor releases, could provide more insight into software variability, whether there are differences between minor releases, and what could be the causes for the observed differences. Importantly, our study does not provide information on analytical accuracy. Reported data shows that there are differences in the estimation of structural metrics between FS versions, however, it is not possible to conclude from our data which version is more accurate.

In conclusion, our results show that the variability across FreeSurfer versions impacts clinical results in the studied PD populations at both subcortical volumetric and vertex-wise levels. The results differed even though we used the same sample as well as the same preprocessing and statistical pipeline. Importantly, estimation of structural metrics is impacted by software variability regardless of the sample with similar results across patients with PD and HC. Observed variability is most likely not reserved solely to FreeSurfer since previous studies reported variability in other analytical software. Our recommendation to users is to facilitate the latest available release of the toolbox unless there is a particular reason to use another release. The most recent software versions usually have improved algorithms and fixed issues discovered in previous releases which ensures staying in line with current standards in the ever-changing field of neuroscience. Software versions should not be updated during studies as it might introduce additional variability to the experiment.

It would be beneficial for the scientific community if developers tracked differences between versions upon release of a new one. This could be achieved by analyzing the same dataset with current and future releases to provide information about the degree of software variability. Developers could also provide long-term support for their software versions, possibly using variability tests such as in Chatelain et al. (2023). Future studies may also implement a multiverse approach to investigate clinical questions with several different software releases.

## Data and Code Availability

The code and results are publicly available at https://github.com/LivingPark-MRI/freesurfer-variability with a notebook detailing the analyses.

## Author Contributions

AS: Conceptualization, Methodology, Software, Investigation, Validation, Formal Analysis, Data Curation, Writing - Original Draft, Visualization. NB: Investigation, Writing - Review & Editing. DK: Writing - Review & Editing, Data Curation. YC: Writing - Review & Editing, Software. MD: Writing - Review & Editing, Software. JBP: Writing - Review & Editing Funding. MS: Writing - Review & Editing, Funding. TG: Conceptualization, Methodology, Investigation, Resources, Supervision, Project Administration, Funding, Writing - Review & Editing.

