## Supplementary material for "The impact of FreeSurfer versions on structural neuroimaging analyses of Parkinson’s disease": Table S1: content.txt

Table S1
Differences between FreeSurfer versions in estimation of subcortical volume across the sample

Table S2
Differences between FreeSurfer versions in estimation of volume, surface area, and cortical thickness across the sample
 
Table S3
Differences between FreeSurfer versions in estimation of volume, surface area, and cortical thickness in healthy controls

Table S4
Differences between FreeSurfer versions in estimation of volume, surface area, and cortical thickness in patients with Parkinson’s disease
 
Table S5
Group differences (healthy controls vs Parkinson’s disease) in FreeSurfer variability 
 
Table S6
Sørensen–Dice coefficients across the sample

Table S7
Differences between FreeSurfer versions in estimation of volume, surface area, and cortical thickness in the replication sample

Table S8
Correlation between quality metrics and software variability in volume across the main sample (FreeSurfer 7 vs 6)

Table S9
Correlation between quality metrics and software variability in surface area across the main sample (FreeSurfer 7 vs 6)

Table S10
Correlation between quality metrics and software variability in cortical thickness across the main sample (FreeSurfer 7 vs 6)

Table S11
Correlation between quality metrics and software variability in volume across the main sample (FreeSurfer 7 vs 5)

Table S12
Correlation between quality metrics and software variability in surface area across the main sample (FreeSurfer 7 vs 5)

Table S13
Correlation between quality metrics and software variability in cortical thickness across the main sample (FreeSurfer 7 vs 5)

Table S14
Correlation between quality metrics and software variability in volume across the main sample (FreeSurfer 6 vs 5)

Table S15
Correlation between quality metrics and software variability in surface area across the main sample (FreeSurfer 6 vs 5)

Table S16
Correlation between quality metrics and software variability in cortical thickness across the main sample (FreeSurfer 6 vs 5)

Table S17
Correlation between quality metrics and software variability in volume across PD in PPMI (FreeSurfer 7 vs 6)

Table S18
Correlation between quality metrics and software variability in surface area across PD in PPMI (FreeSurfer 7 vs 6)

Table S19
Correlation between quality metrics and software variability in cortical thickness across PD in PPMI (FreeSurfer 7 vs 6)

Table S20
Correlation between quality metrics and software variability in volume across PD in PPMI (FreeSurfer 7 vs 5)

Table S21
Correlation between quality metrics and software variability in surface area across PD in PPMI (FreeSurfer 7 vs 5)

Table S22
Correlation between quality metrics and software variability in cortical thickness across PD in PPMI (FreeSurfer 7 vs 5)

Table S23
Correlation between quality metrics and software variability in volume across PD in PPMI (FreeSurfer 6 vs 5)

Table S24
Correlation between quality metrics and software variability in surface area across PD in PPMI (FreeSurfer 6 vs 5)

Table S25
Correlation between quality metrics and software variability in cortical thickness across PD in PPMI (FreeSurfer 6 vs 5)

Table S26
Correlation between quality metrics and software variability in volume across PD in QPN (FreeSurfer 7 vs 6)

Table S27
Correlation between quality metrics and software variability in surface area across PD in QPN (FreeSurfer 7 vs 6)

Table S28
Correlation between quality metrics and software variability in cortical thickness across PD in QPN (FreeSurfer 7 vs 6)

Table S29
Correlation between quality metrics and software variability in volume across PD in QPN (FreeSurfer 7 vs 5)

Table S30
Correlation between quality metrics and software variability in surface area across PD in QPN (FreeSurfer 7 vs 5)

Table S31
Correlation between quality metrics and software variability in cortical thickness across PD in QPN (FreeSurfer 7 vs 5)

Table S32
Correlation between quality metrics and software variability in volume across PD in QPN (FreeSurfer 6 vs 5)

Table S33
Correlation between quality metrics and software variability in surface area across PD in QPN (FreeSurfer 6 vs 5)

Table S34
Correlation between quality metrics and software variability in cortical thickness across PD in QPN (FreeSurfer 6 vs 5)


Figure S1. Estimation of volume of subcortical regions in Freesurfer 5, 6, and 7.
 
Figure S2. Estimation of volume of cortical regions in Freesurfer 5, 6, and 7.
 
Figure S3. Estimation of surface areas in Freesurfer 5, 6, and 7.
 
Figure S4. Estimation of cortical thickness in Freesurfer 5, 6, and 7.

Figure S5. The volume of the left hippocampus. (A) between-version variability in the same subject. FreeSurfer 7 in red,FreeSurfer 5 in blue, overlap in pink. (B) between-subject variability in the same FreeSurfer version 7. Subject 1 in red, subject 2 in blue, overlap in pink. Image from subject 2 registered to subject 1 with rigid transformation (6 degrees of freedom). I, inferior view; L, left view; R, right view; S, superior view.
