## Supplementary figures and images for "The impact of FreeSurfer versions on structural neuroimaging analyses of Parkinson’s disease"

### Fig_S1.png

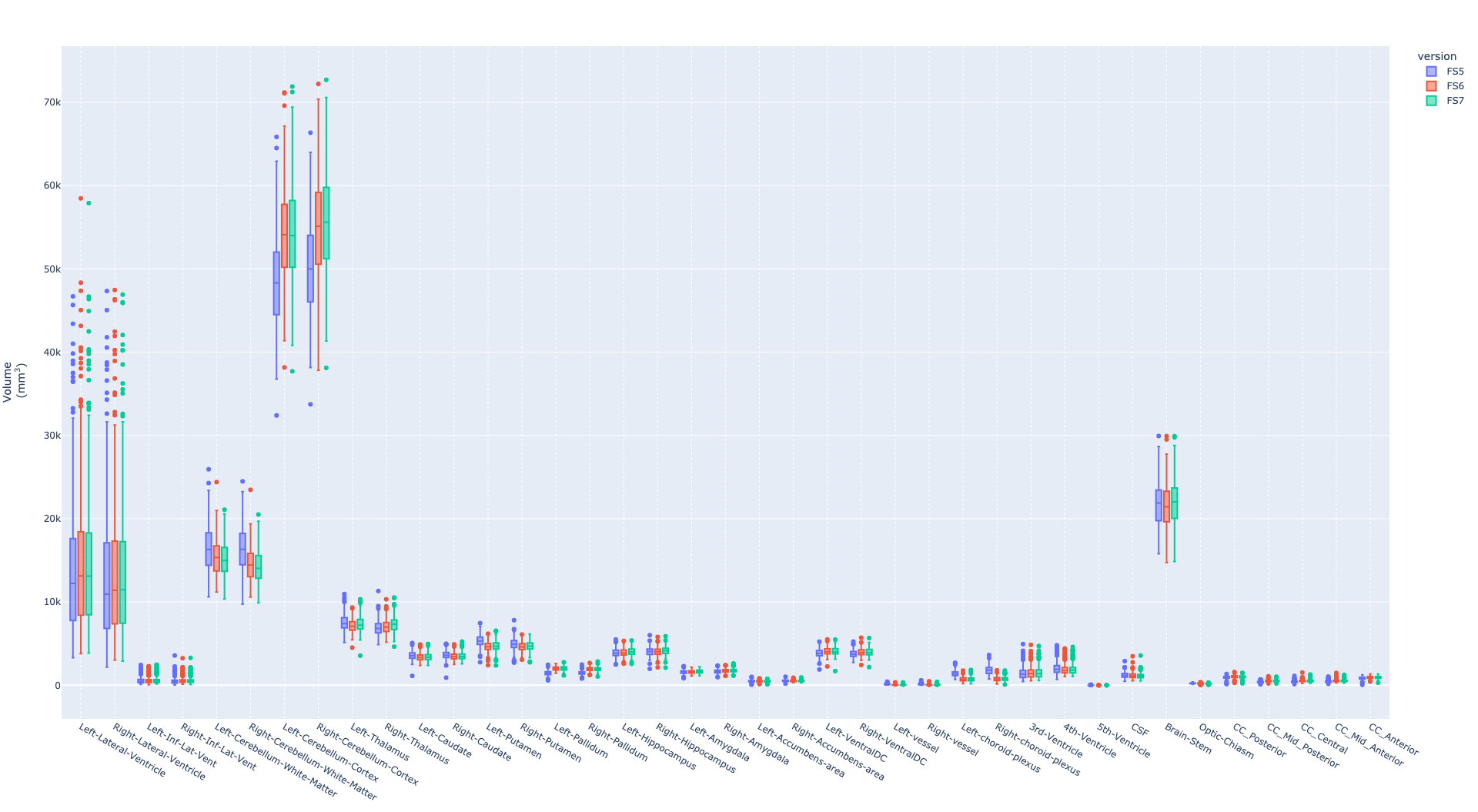

### Fig_S2.png

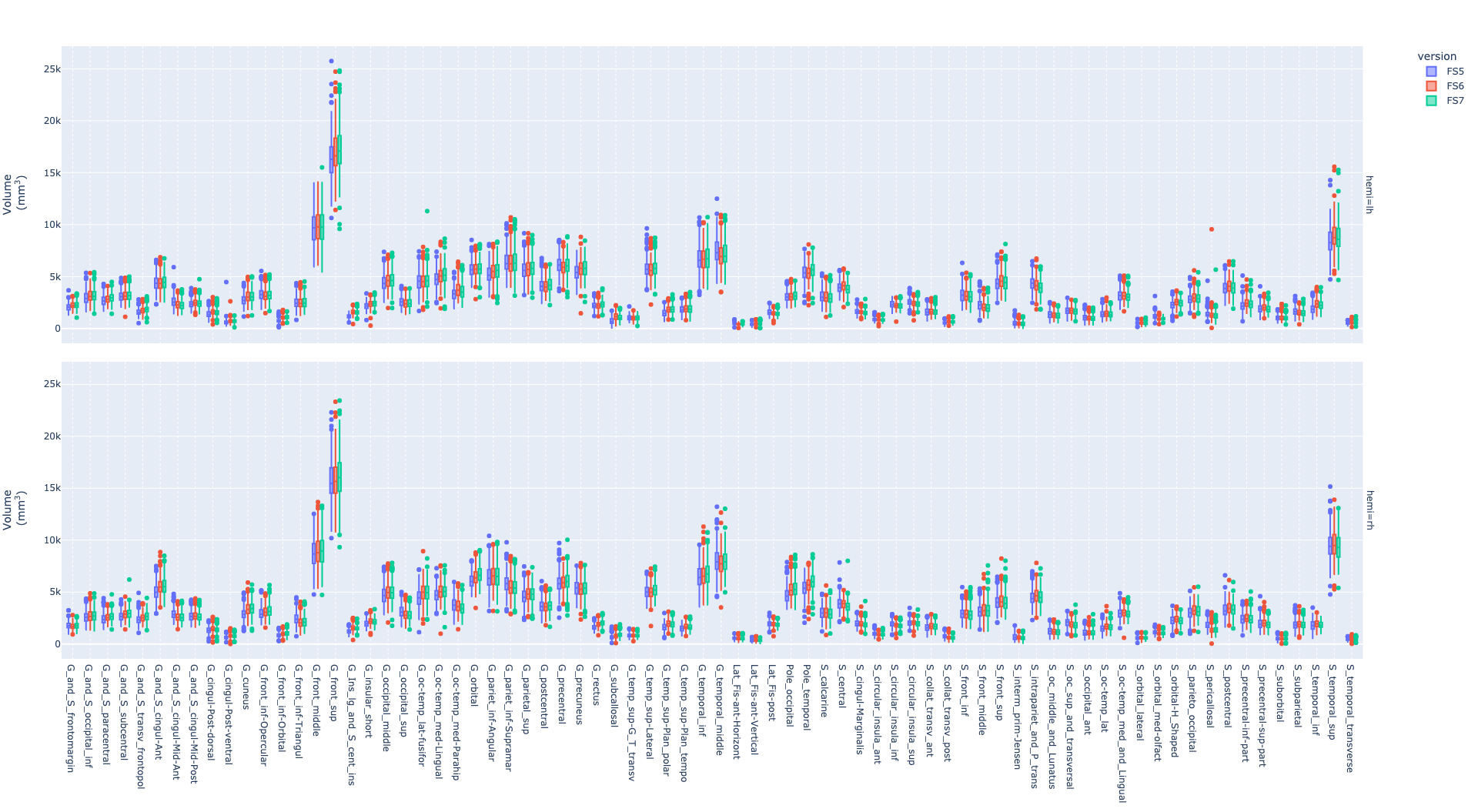

### Fig_S3.png

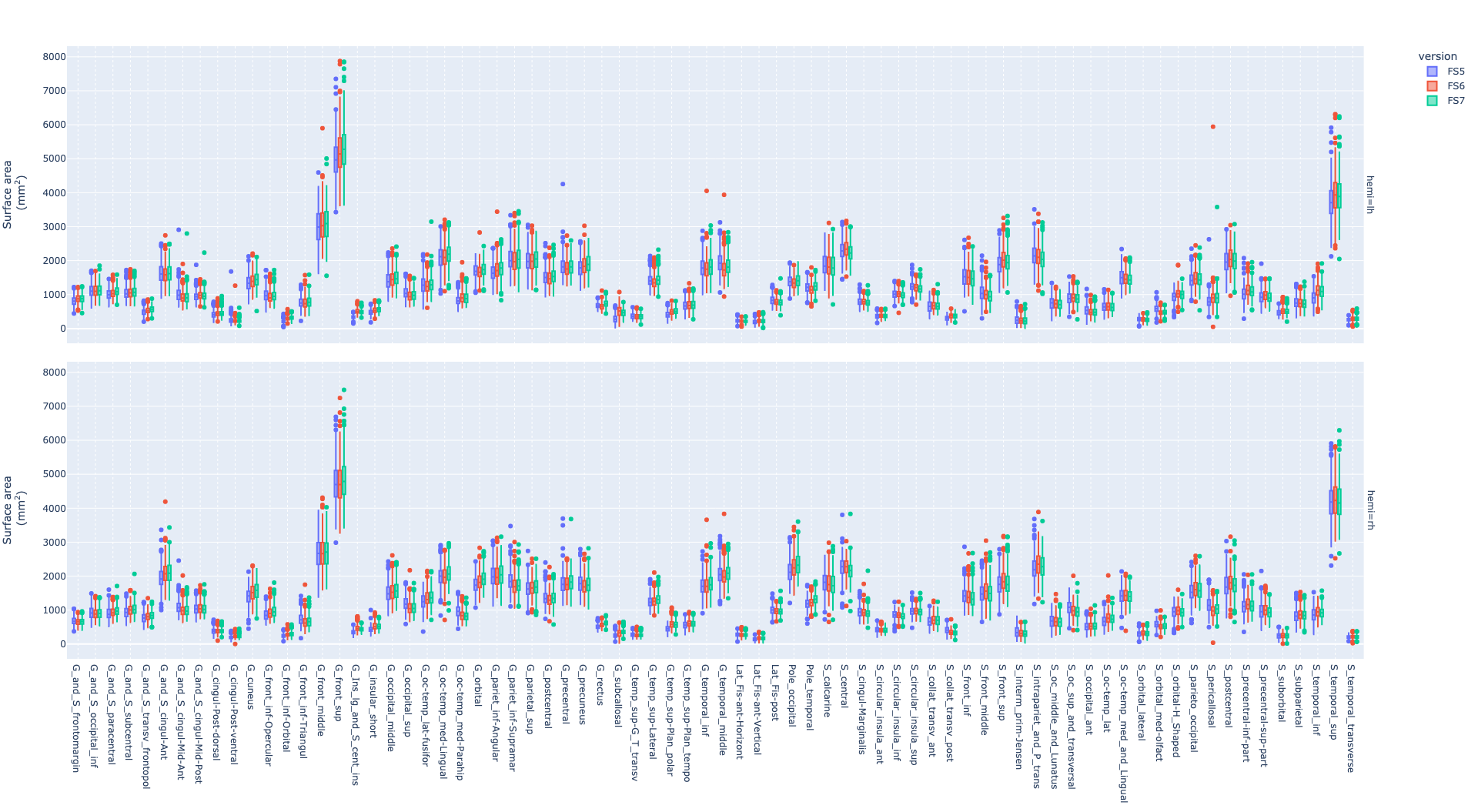

### Fig_S4.png

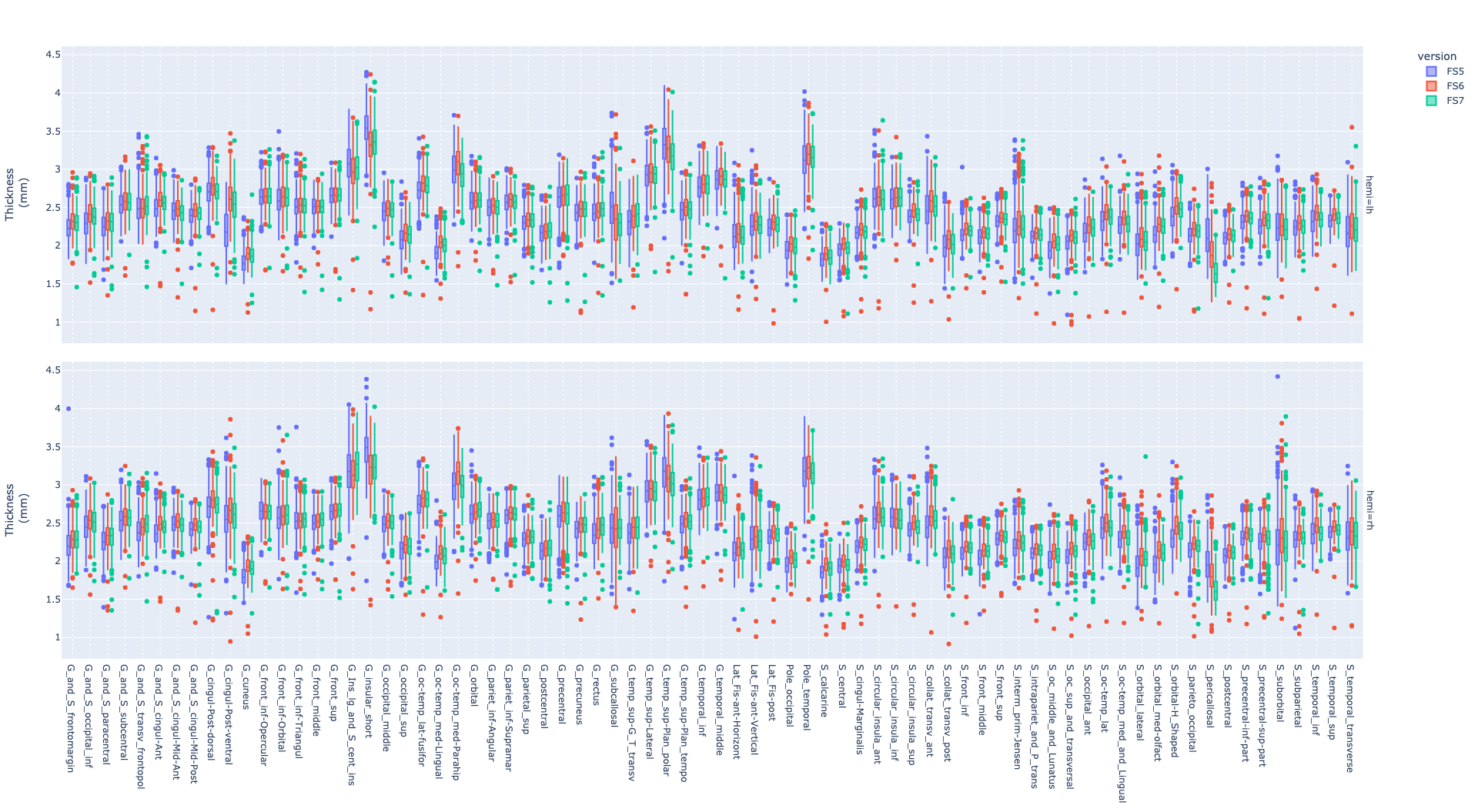

### Fig_S5.png

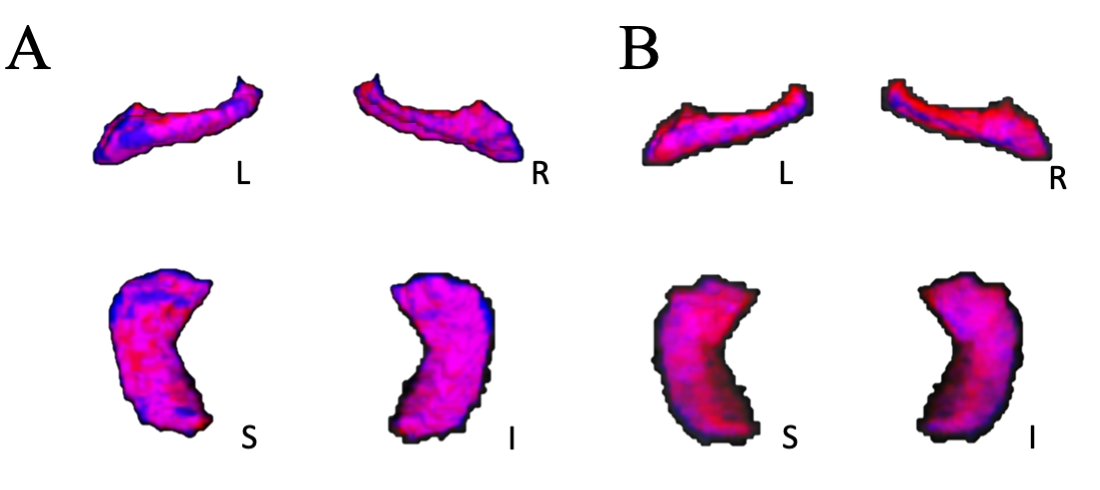

### Fig_S6.png

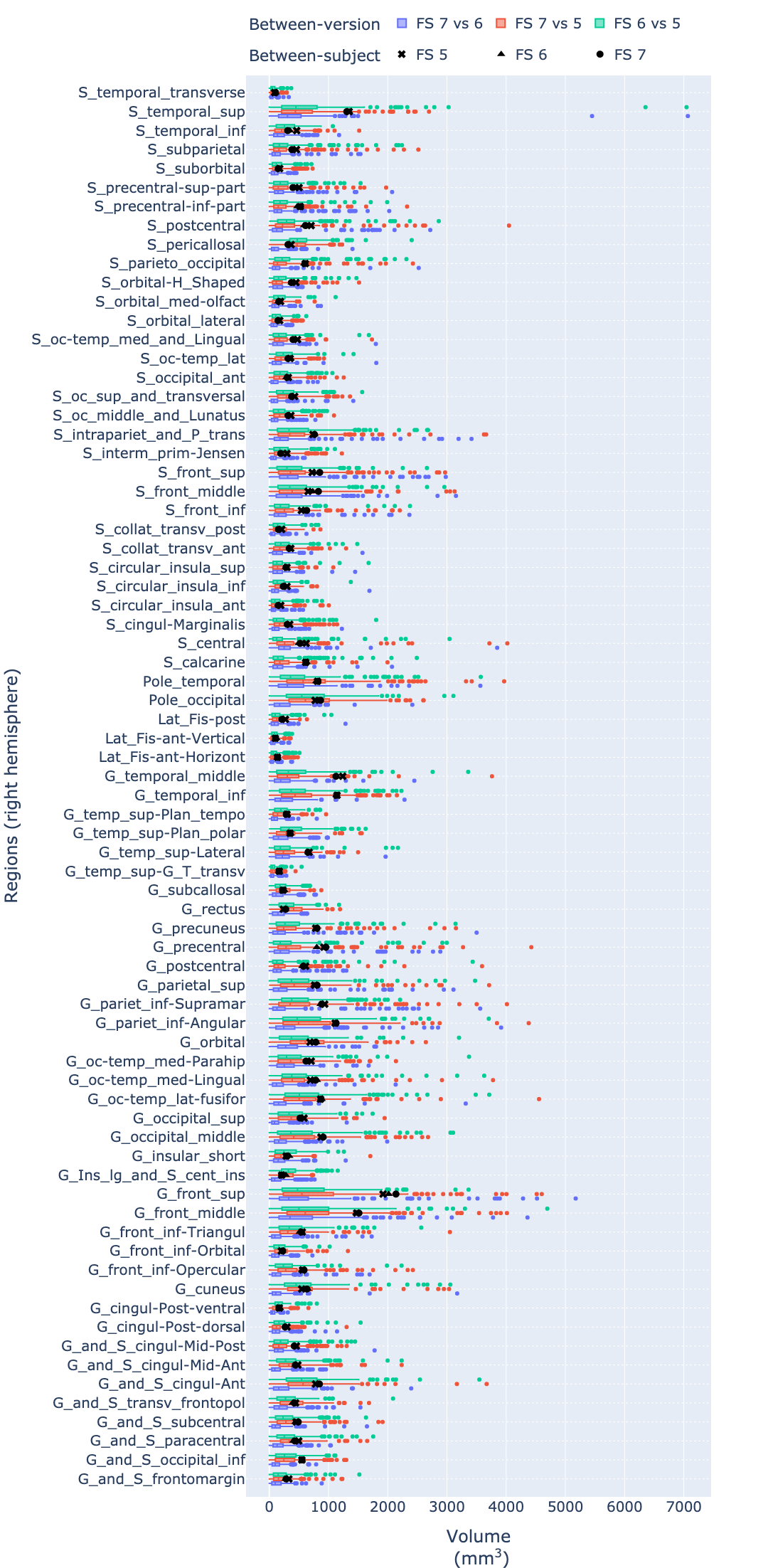

### Fig_S7.png

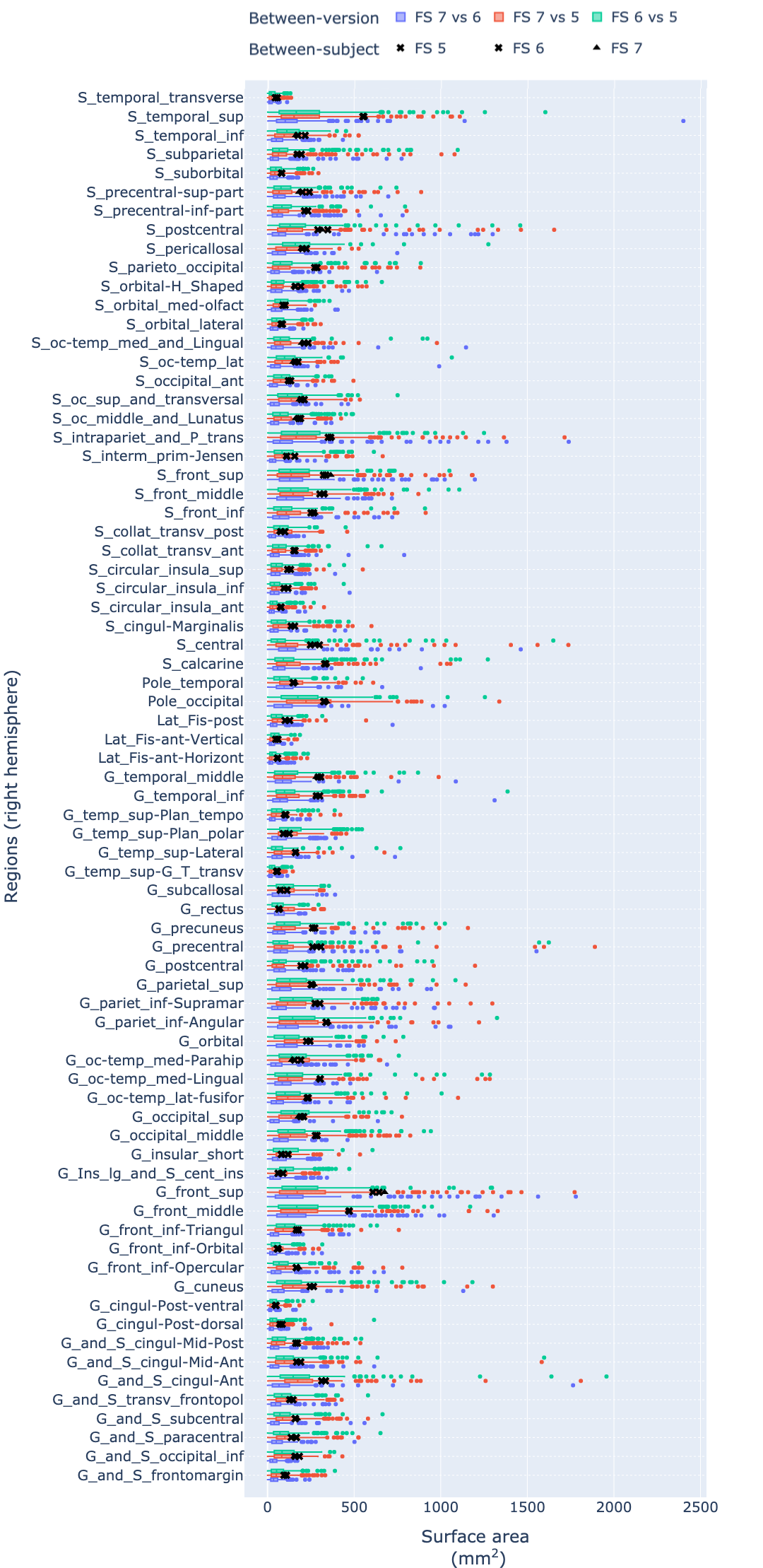

### Fig_S8.png

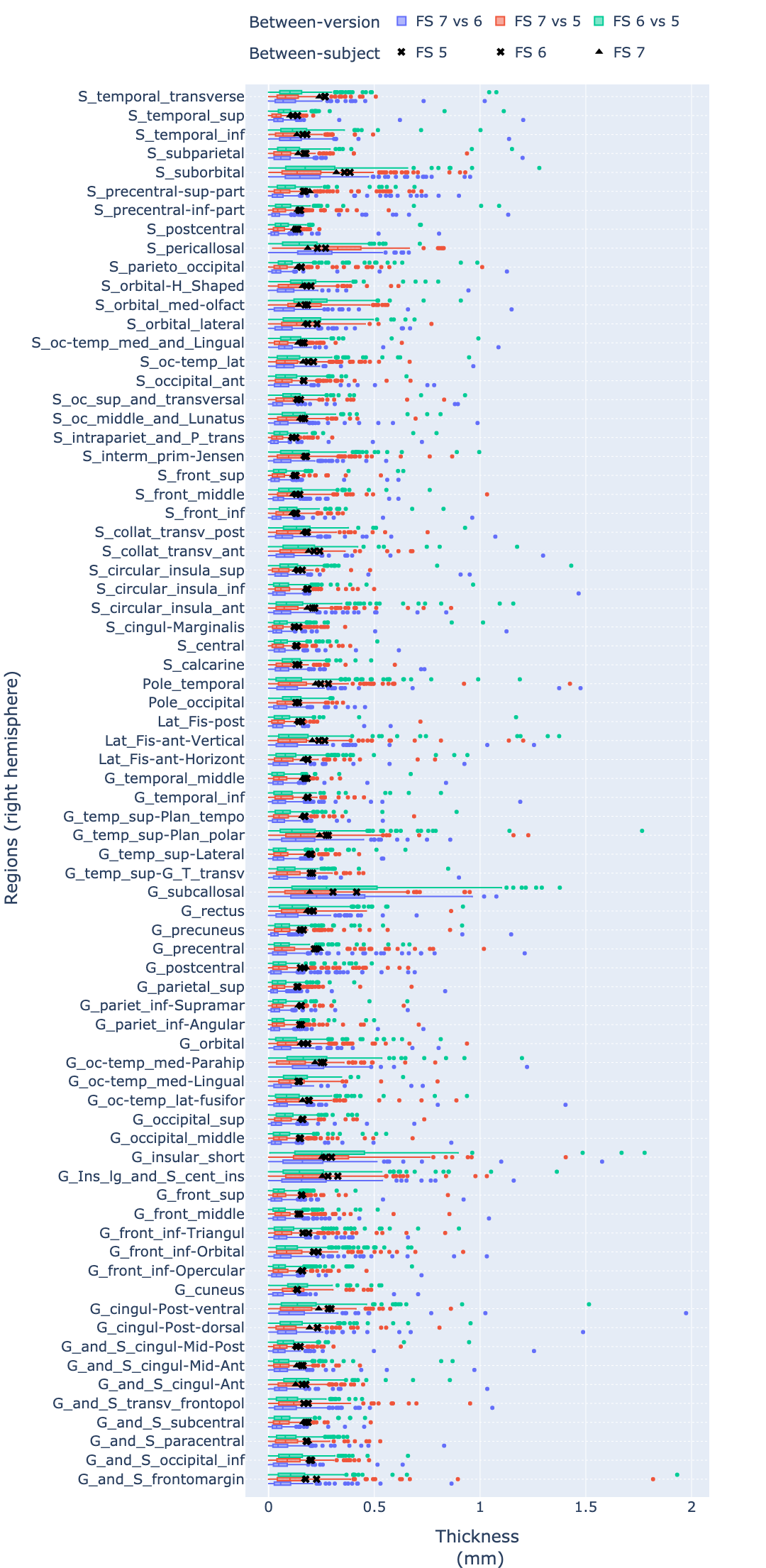
